# Nuclear pores constrict upon energy depletion

**DOI:** 10.1101/2020.07.30.228585

**Authors:** Christian E. Zimmerli, Matteo Allegretti, Vasileios Rantos, Sara K. Goetz, Agnieszka Obarska-Kosinska, Ievgeniia Zagoriy, Aliaksandr Halavatyi, Julia Mahamid, Jan Kosinski, Martin Beck

## Abstract

Nuclear pore complexes (NPCs) fuse the inner and outer nuclear membranes and mediate nucleocytoplasmic exchange. They are made of 30 different nucleoporins that form an intricate cylindrical architecture around an aqueous central channel. This architecture is highly dynamic in space and time. Variations in NPC diameter were reported, but the physiological circumstances and the molecular details remain unknown. Here we combined cryo-electron tomography and subtomogram averaging with integrative structural modeling to capture a molecular movie of the respective large-scale conformational changes *in cellulo*. While actively transporting NPCs adopt a dilated conformation, they strongly constrict upon cellular energy depletion. Fluorescence recovery after photo bleaching experiments show that NPC constriction is concomitant with reduced diffusion and active transport across the nuclear envelope. Our data point to a model where the energy status of cells is linked to the conformation of NPC architecture.

## Introduction

Nuclear pore complexes (NPCs) bridge the nuclear envelope (NE) and facilitate nucleocytoplasmic transport. Across the eukaryotic kingdom, about 30 different genes encode for NPC components, termed nucleoporins (Nups). Although specialized Nups have been identified in many species, extensive biochemical and structural studies *in vitro* led to the consensus that the core scaffold inventory is conserved. It consists of several Nup subcomplexes that come together in multiple copies to form an assembly of eight asymmetric units, called spokes, that are arranged in a rotationally symmetric fashion (*1*). The Y-complex (also called Nup107 complex) is the major component of the outer rings (the nuclear and cytoplasmic rings; NR and CR), which are placed distally into the nuclear and cytoplasmic compartments. The inner ring complex scaffolds the inner ring (IR; also called spoke ring) that resides at the fusion plane of the nuclear membranes. The Nup159 complex (also called P-complex) asymmetrically associates with the Y-complex of the cytoplasmic ring and mediates mRNA export. Despite these common features of quaternary structure, *in situ* structural biology studies have revealed that the higher order assembly is variable across the eukaryotic kingdom (*2*).

In addition to compositional variability across different species, NPC architecture is conformationally highly dynamic and variations in NPC diameter have been observed in various species and using different methods (*3*–*7*). It has been shown that dilation is part of the NPC assembly process (*8*, *9*). However, if NPC dilation and constriction may play a role during active nuclear transport (*10*), or are required to open up peripheral channels for the import of inner nuclear membrane proteins (*11*–*13*), remains controversial. It is difficult to conceive that such large-scale conformational changes can occur on similar time scales as individual transport events (*14*, *15*), which would be the essence of a physical gate. Nevertheless, several cues that potentially could affect NPC diameter have been suggested, such as exposure to mechanical NE stress, mutated forms of Importin β, varying Ca^2+^ concentrations or hexanediol (*7*, *16*–*21*). However, these previous studies did neither explore NPC diameter and its functional consequences within intact cellular environments nor did they structurally analyze the conformational changes of nuclear pores in molecular detail. Thus, physiological cause and consequence along with the molecular mechanisms of NPC dilation and constriction remain enigmatic.

Active nuclear transport of cargo relies on energy supply. Importin or exportin-mediated transport requires the small GTPase RAN that binds either GTP in the nucleus or GDP in the cytoplasm to ensure directionality of transport (*22*), while mRNA export is directly ATP dependent (*23*). Cells of various organisms including *Schizosaccharomyces pombe* show a rapid shut down of active nuclear transport and mRNA export when depleted of ATP (*24*–*26*). This points to a well conserved mechanism, likely dependent on a concomitantly reduced availability of free GTP (*27*). Moreover, energy depletion (ED) leads to a general reorganization of the cytoplasm including solidification of the periplasm, general water loss and reduction of the nuclear and cellular volume, which allows cells to endure under unfavorable conditions (*28*–*31*). If the shutdown of active nuclear transport coincides with the alteration in passive diffusion and potentially a conformational adaption of NPC architecture remains unknown.

Here we demonstrate that in *S. pombe* NPCs (SpNPCs) constrict under conditions of ED, which is concomitant with a reduction of both, free diffusion and active nuclear transport across the nuclear envelope. Using *in cellulo* cryo-electron microscopy (cryo-EM) and integrative structural modeling, we captured a molecular movie of NPC constriction. Our dynamic structural model suggests large scale conformational changes that occur by movements of the spokes with respect to each other but largely preserve the arrangement of individual subcomplexes. Previous structural models obtained from isolated nuclear envelopes (*32*–*37*) thereby represent the most constricted NPC state.

### *In cellulo* cryo-EM map of the *S. pombe* NPC

To study NPC architecture and function *in cellulo* at the best possible resolution and structural preservation, we explored various genetically tractable model organisms for their compatibility with cryo-focused ion beam (FIB) specimen thinning, cryo-electron tomography and subtomogram averaging (STA). *Saccharomyces cerevisiae* cells were compatible with high throughput generation of cryo-lamellae and acquisition of tomograms. STA of their NPCs resulted in moderately resolved structures (*4*). In contrast, a larger set of cryo-tomograms from *Chaetomium thermophilum* cells did not yield any meaningful averages, possibly because their NPCs displayed a very large structural variability. We therefore chose to work with *S. pombe* cells that are small enough for thorough vitrification, offer a superior geometry for FIB-milling compared to *C. thermophilum* with the advantage of covering multiple cells and, compared to *S. cerevisiae*, higher number of NEs and NPCs per individual cryo-lamellae and tomogram, leading to high data throughput (**Fig. S1**).

To obtain a high quality cryo-EM map of *S. pombe* NPCs, we prepared cryo-FIB milled lamellae of exponentially growing *S. pombe* cells and acquired 178 tomograms from which we extracted 726 NPCs. Subsequent STA resulted in an *in cellulo* NPC average of very high quality in terms of both visible features (**Fig. 1A, B and Fig. S2A**) and resolution (**Fig. S2B**). Systematic fitting of the *S. pombe* IR asymmetric unit model (see Materials and Methods), resulted in a highly significant fit (**Fig S3A**). The subsequent refinement with integrative modeling led to a structural model that explains the vast majority of the observed electron optical density in the IR (**Fig. 1B**, **Fig. S4, and Video S1**). The IR architecture appears reminiscent to NPC structures of other eukaryotes (**Fig. S5**) further corroborating its evolutionary conservation (*1*).

**Figure 1.**
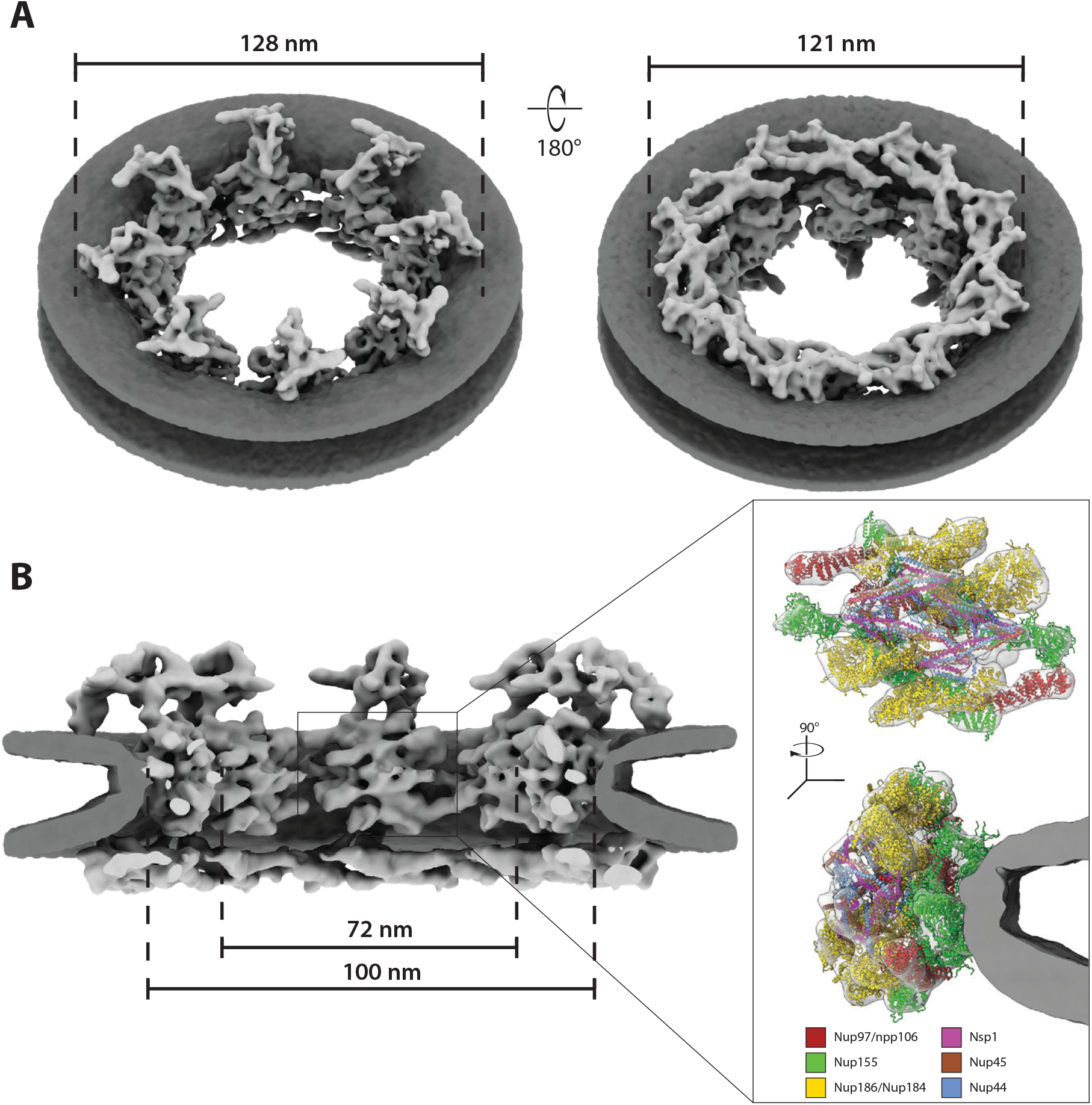
*In cellulo* cryo-EM map of the S. pombe NPC. A) Isosurface rendered views of the *S. pombe* NPC as seen from the cytoplasm (left) and the nucleoplasm (right; with membranes in dark and protein in light grey). While the cytoplasmic view (left) reveals eight disconnected protein entities instead of a cytoplasmic ring, the nuclear view (right) shows two concentric nuclear Y-complex rings. B) Same as (A) but shown as cutaway view. While the asymmetric curvature of the nuclear membranes and the arrangement of the cytoplasmic side is unprecedented in other species, the inner ring architecture is highly conserved as highlighted in the inset (see also **Fig. S5**). Fitting of inner ring nucleoporin homology models explains the vast majority of the observed electron optical density.

Although the outer rings are known to be more variable, the intra-subcomplex interaction network of the Y-complex (Nup120, Nup85, Nup145C, Sec13, Nup84 and Nup133) has been comprehensively characterized by many studies and considered to be conserved (*1*) (see **Table S1** for nomenclature of Nups across different species). Systematic fitting revealed that the NR of the SpNPC is composed of two concentric Y-complex rings (**Fig. 1A**, **Fig. S3B and Video S1**) as in vertebrates and algae but as opposed to the single Y-complex ring observed in *S. cerevisiae* (**Fig. S2A**) (*4*, *35*, *38*). Integrative modeling of the entire Y-complex ensemble of the NR revealed a rather classical Y-complex architecture with the typical head-to-tail oligomerization (**Fig. 2A and Fig. S4**). This analysis emphasizes that *S. pombe* Y-complex Nups do localize to the NR, contrasting previous proposals (*39*). The homology models of SpNup131 and SpNup132 fit to the Y-complex tail region equally well, rendering these two proteins indistinguishable by our approach.

**Figure 2.**
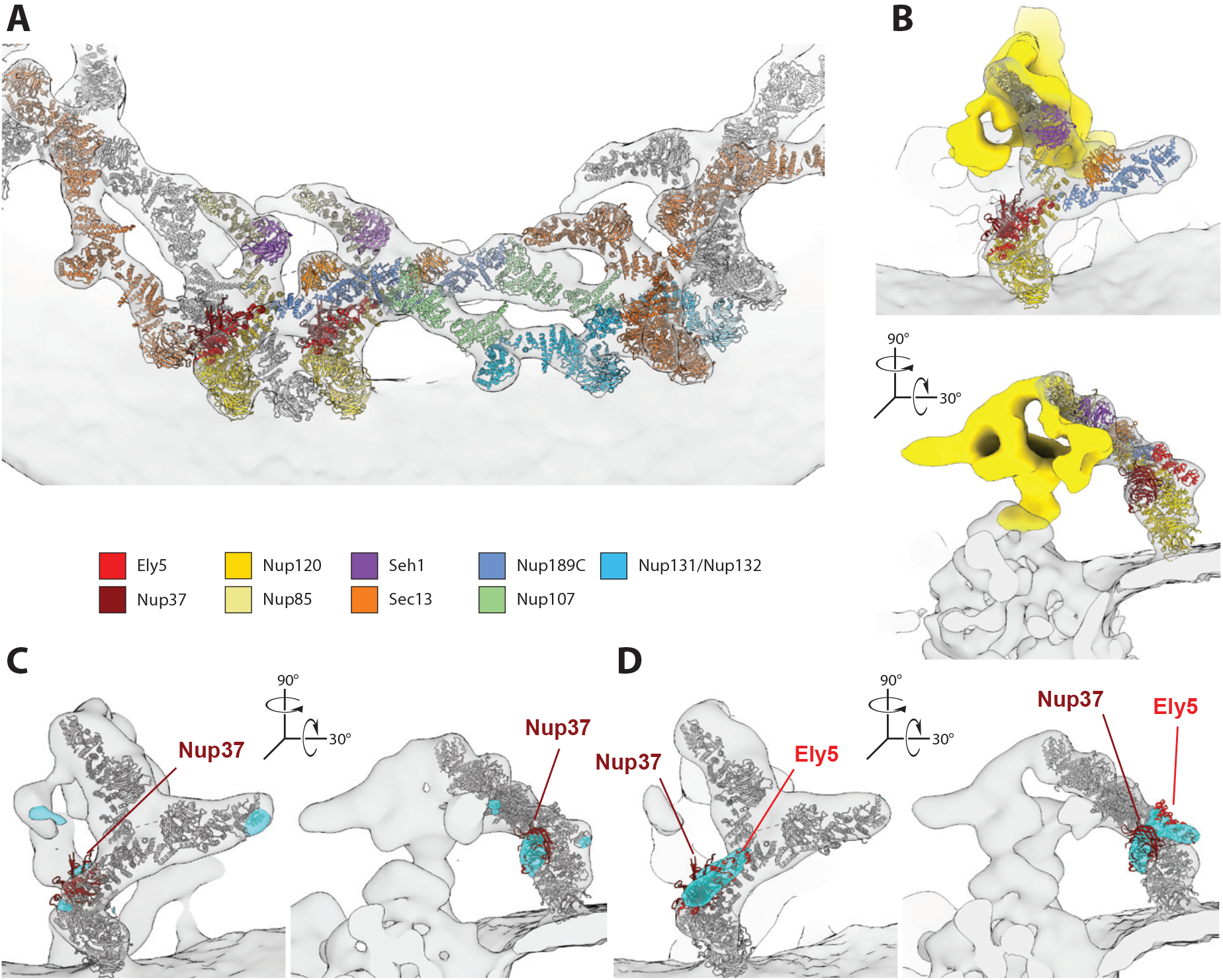
Architecture of spNPC outer rings. A) Systematic fitting and integrative modeling of all *S. pombe* Y-complex nucleoporins reveals a head-to-tail arrangement with two concentric Y-complex rings on the nuclear side of the SpNPC similar to humans. A segment of the NR of the cryo-EM map is shown isosurface rendered in transparent light grey. The adjacent inner Y-complexes are shown in grey and outer Y-complexes are shown in orange. B) Integrative model of the cytoplasmic protein entities. The fit of the Y-complex vertex explains most of the observed density. The mRNA export platform as identified in (*4*, *62*) is segmented in yellow. C, D) Verification of the molecular identity of the observed structure. C) The *nup37Δ* cryo-EM map is shown in light grey and overlaid with the difference map (cyan) of the *wild type* and *nup37Δ* maps, both filtered to 27 Å. The missing density in the long arm of the Y-vertex coincidences with the position of Nup37 (dark red) of the Y-complex vertex (dark grey, as in A). D) *nup37Δ*-*ely5Δ* double knockout map (light grey) overlaid with the corresponding difference map (cyan). Differences are apparent at the location of both, Nup37 (dark red) and Ely5 (light red) with respect to the fitted Y-complex model (dark grey).

Closer inspection of the cytoplasmic side of cryo-EM map revealed a surprising and unprecedented architectural outline, since it did not form a ring. Instead, eight spatially separated entities were observed (**Fig. 1A**) suggesting that the integrity of the cylindrical outline is rather provided by the IR and NR while the cytoplasmic protein entities serve as a mere anchor point for the mRNA export platform, the Nup159 complex (**Fig. 2B**). Although the dynein-arm that is characteristic for the *S. cerevisiae* NPC (*40*) is lacking, the Nup159 complex resembles its *S. cerevisiae* counterpart in shape (**Fig. S6A**). Systematic fitting and subsequent refinement with integrative modeling revealed that the Y-complex vertex fits into the density observed at the cytoplasmic side (**Fig. 2B**, **Fig. S3C, Fig. S4 and Video S1**). The density potentially accounting for Nup107 and SpNup131/SpNup132 was missing (**Fig. S3D**) and could not be recovered by local refinement (**Fig. S6B**). Instead, the observed density sharply declined at the edge of SpNup189C consistent with previous work suggesting a split of the SpNup189C-Nup107 interface (**Fig. S6C**) (*39*, *41*). To independently confirm the identity of the observed vertex-like density, we analyzed *nup37Δ* and *nup37Δ*-*ely5Δ* strains in which non-essential, peripheral Y-complex Nups were deleted. The binding of both, Nup37 and Ely5 to Nup120 has been previously shown *in vitro* (*42*, *43*), and as expected, density was missing in the respective positions of all Y-complexes (**Fig. 2C-D and Fig. S7A-B**). Unexpectedly, a density that could accommodate Ely5 homology model was missing also in the cytoplasmic Y-complex, suggesting that Ely5 is present in *S. pombe* at both, the nuclear and the cytoplasmic side of the NPC (**Fig 2D and Video S1**) unlike in higher metazoans where its homolog ELYS is known to exclusively bind to the NR (*35*, *44*). Otherwise, the NPC architecture remained mostly unchanged, despite some increased flexibility in the Nup120 arm of the outer nuclear Y-complex (**Fig. S7A-B**). These results unambiguously identify the density observed at the cytoplasmic side as *bona fide* Y-complex vertex.

### Energy depletion leads to constriction of NPC scaffold and central channel

Previous cryo-EM structures of NPCs obtained from isolated nuclear envelopes (*32*–*37*) or by detergent extraction (*45*) had a smaller diameter as compared to those obtained from intact cells (*3*, *4*, *38*, *46*). We therefore hypothesized that NPC diameter may depend on the biochemical energy level that is depleted during preparations of isolated nuclear envelopes or NPCs but may also be reduced within intact cells e.g. during stress conditions. We set out to systematically analyze the NPC architecture under conditions of energy depletion as compared to exponentially growing cells. We structurally analyzed 292 NPCs subsequent to 1 hour of ED using non-hydrolysable 2-deoxy-glucose in combination with the respiratory chain inhibitor antimycin A (see Materials and Methods). We measured the diameter based on centroids of the spokes as obtained by STA (see Materials and Methods) and found a significant constriction of the mean central channel diameter during ED from ~70 nm to ~55 nm (**Fig. 3A**). The variation of diameters was larger within the population of NPCs exposed to ED as compared to the actively transporting conditions, which likely blurred structural features during STA. To generate a conformationally more homogenous ensemble, we split the particles form the ED data set into two classes with central channel diameters of <50 nm and >50 nm (533 and 1012 subunits respectively) (**Fig. 3B**) and refined them separately to <28 Å resolution (**Fig. 3C-D and Fig. S8A-B**). Both conformations of the ED state showed a smaller NPC diameter compared to the control. The intermediate conformation was ~65 nm wide at the IR while the most constricted conformation showed a diameter of ~49 nm (**Fig. 3D**) and is thus comparable to the diameter observed in isolated NEs (*32*–*37*). We further calculated the diameters at the level of the cytoplasmic side and NR and found that all three rings constrict significantly during ED. While the diameter of the IR and cytoplasmic side changed their conformation most dramatically, the NR was less affected (**Fig. 3E**, **Videos S4-S6**). The estimated volume of the central channel in the most dilated state was almost twice as large (~152’000 nm^3^) as compared to the most constricted state (~86’000 nm^3^) (**Fig. S8C**), which likely translates to ~2-fold change in concentration of the FG-repeats contained therein.

**Figure 3.**
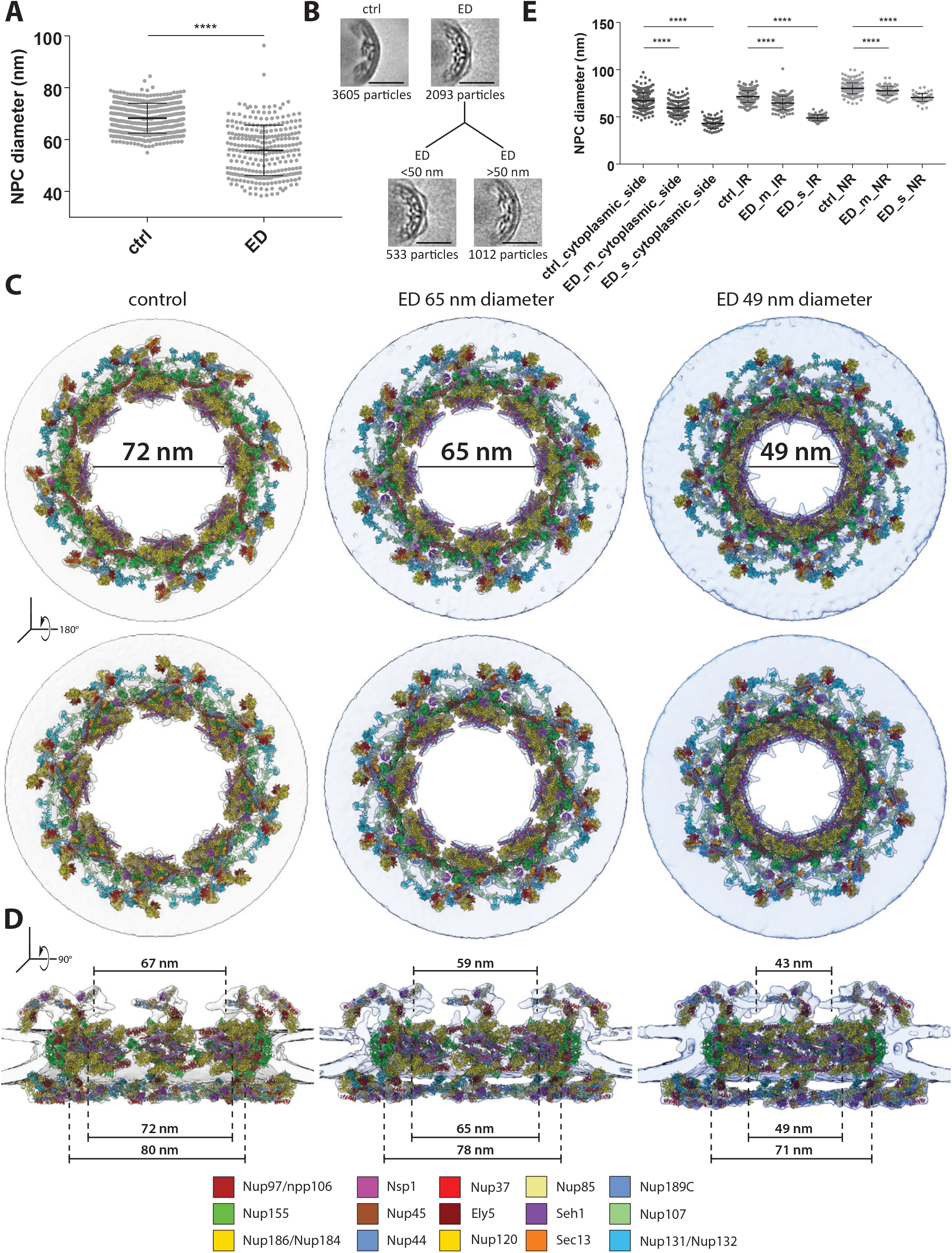
NPCs constrict upon energy depletion. A) Diameter measurement of control (n=438) and energy depleted (n=271) NPCs based on subunit positions obtained by STA (see Materials and methods) reveals a significant constriction of NPCs upon ED (mean ± standard error of mean (SdEM) control: 68.2 nm and mean ED: 55.84 nm, whiskers indicate standard deviation, p-value < 0.0001, two-sided t-test). B) Slices through subtomogram averages corresponding to (A) show the different conformations at the level of the IR. NPCs were divided into two classes with diameters of <50 nm (left) and >50 nm (right; scale bar: 50 nm). C) Measurements of individual NPC diameters at the CR, IR and NR in control conditions and during ED reveal significant diameter constriction of all three rings (all p-values <0.0001; ctrl control; ED_m represents the class with diameter of >50nm; ED_s represents the class with diameter of <50 nm. Diameters mean ± SdEM measured are: 67.25 ± 0.44, n=341 (ctrl_cytoplasmic_side); 59.29 ± 0.6817, n=136 (ED_m_cytoplasmic_side) and 43.15 ± 0.6509, n=68 (ED_s_cytoplasmic_side); 71.59 ± 0.3459, n=341 (ctrl_IR); 64.73 ± 0.626, n=136 (ED_m_IR) and 49 ± 0.3815, n=68 (ED_s_IR); 80.22 ± 0.3106, n=341 (ctrl_NR); 77.71 ± 0.4006, n=136 (ED_m_NR) and 70.59 ± 0.5001, n=68, whiskers indicate standard deviation. D) Cytoplasmic view (top) and nuclear view (bottom) of cryo-EM maps superimposed with the respective integrative models from actively transporting, intermediate and fully constricted NPCs illustrating the overall conformational change leading to a central channel diameter constriction from 72 nm to 49 nm. The cytoplasmic and IR spokes move as individual entities and contribute the most to the central channel diameter change, whereas the NR constricts to a lesser extent. E) Same as (D) but show as cutaway side view. Upon constriction, the mRNA export platform bends towards the center of the NPC. Conformational changes of the NR are less dramatic and include mostly changes in the curvature of the Y complexes (see also supplementary videos 4-8).

To better understand how NPCs accommodate such massive conformational changes on the molecular level, we systematically fitted individual subcomplexes (**Fig. S9**) and built structural models of the three different diameter states based on the cryo-EM maps (**Fig. 3C-D)** using a multi-state integrative modeling procedure (**Fig S4**). In the cytoplasmic side and the NR, conformational changes were limited to the curvature of the Y-complexes and inward-bending of the mRNA export platform towards the center of the pore (**Videos S4-6**). In contrast, the central channel constriction of the IR is more elaborate and mediated by a lateral displacement of the 8 spokes that move as independent entities to constrict or dilate the IR (**Fig. 3D and Videos S4-S6**). In the dilated state, around 3-4 nm wide gaps are formed in-between the neighboring spokes, while in the constricted state the spokes form extensive contacts (**Fig. S10A-B**), equivalent to those in the previously published structures of the human NPC in isolated nuclear envelopes (*36*). Notably, the spokes do not move entirely as rigid bodies, but some conformational changes occur within the Nup155 and Nsp1 complex regions (**Video S7 and S8**). Those are however distinct from the previously proposed conformational sliding (*10*) and rather consistent with an overall preserved intra-subcomplex arrangement (*15*, *47*).

Interestingly, under conditions of ED additional density is arching out into the lumen of the NE (**Fig S10A and C**), contrasting control conditions under which they are less clearly discernible from the membrane. It has been previously proposed that such luminal structures are formed by Pom152 (ScPom152 or HsGP210) (*48*, *49*). In terms of their shape, the observed arches are reminiscent to those observed in isolated Xenopus *laevis* (*34*). Our data imply that the luminal ring conformation becomes more prominent upon constriction. If this has any mechanical benefits to keep NPCs separated (*34*), or might rather limit the maximal dilation, remains to be further investigated.

### NPC constriction is concomitant with reduced diffusion and active nuclear transport

We wondered about the transport competence of NPCs in conditions under which they are constricted in comparison to actively transporting, dilated NPCs. To address this, we employed live cell imaging of *S. pombe* cells expressing a GFP variant tagged with a nuclear localization signal (NLS) on its N- and C-terminus (NLS-GFP) that shows a nuclear localization under control conditions (**Fig. 4A**). Already after 30 min of ED most of the NLS-GFP localized into the cytoplasm (**Fig. 4 B**), confirming that active nuclear import is suspended (*26*). To assess passive diffusion across the nuclear envelope, we performed fluorescence recovery after photobleaching (FRAP) experiments of nuclei in cells expressing freely diffusing GFP at different time points after ED as compared to control conditions (**Fig. 4C and Fig. S11A-B**) (see Materials and Methods). GFP diffusion rates into the nucleus were significantly decreased upon energy depletion (**Fig. 4C**), contrasting a minor, negligible effect observed within the cytoplasm (**Fig. S11C**). Passive diffusion was the slowest after about 1 hour of ED, the time at which we structurally analyzed NPC architecture and is thus concomitant with NPC constriction (**Fig. 4C**).

**Figure 4.**
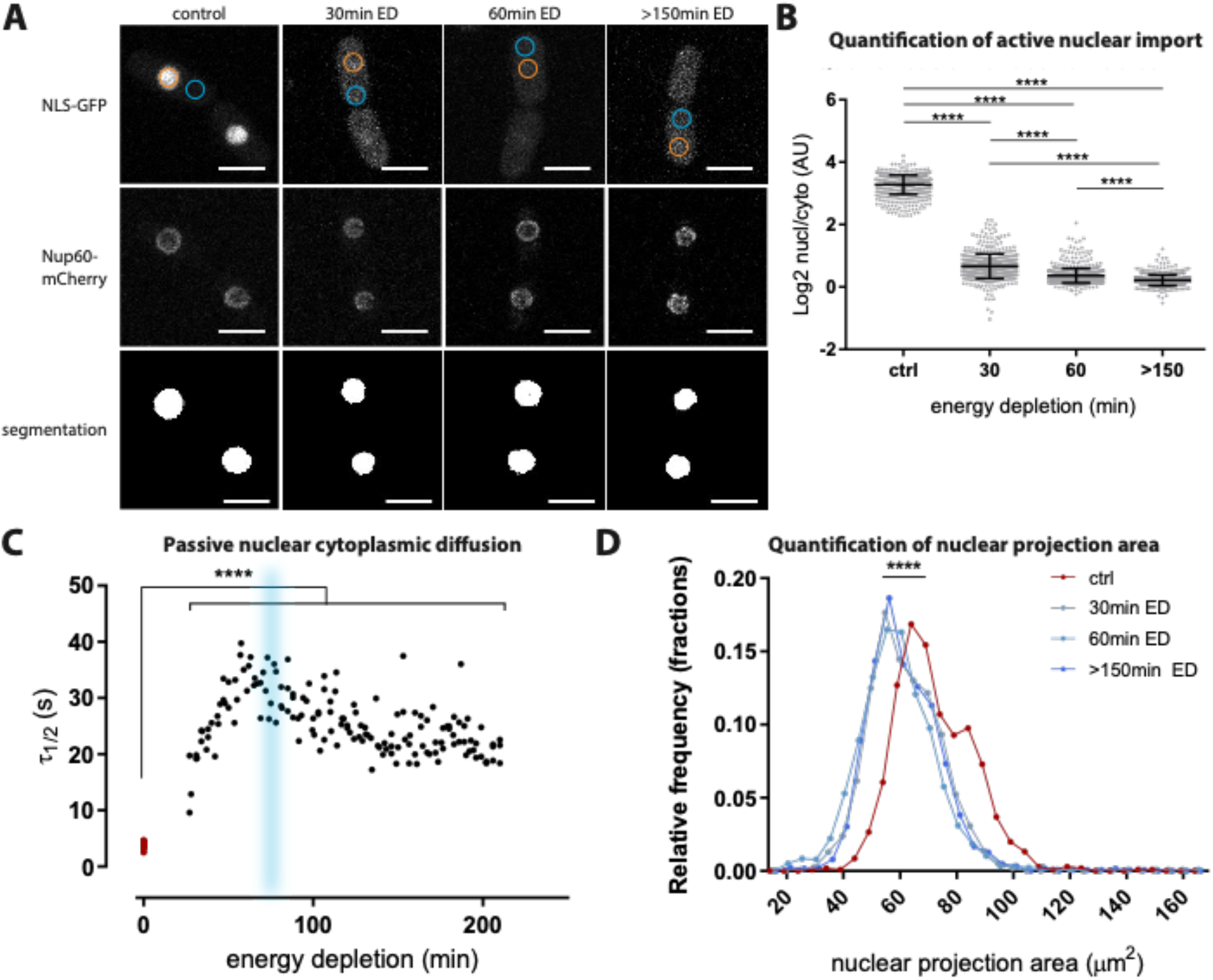
How energy depletion affects transport across the nuclear envelope and nuclear size. A) Maximum projected confocal stack from life cell imaging of NLS-GFP and Nup60-mCherry are used to measure active nuclear transport and nuclear size during energy depletion. Actively imported NLS-GFP loses its nuclear localization during ED indicating a shutdown of active nuclear transport during ED. The nuclear projection area is reduced during ED as determined by segmenting the Nup60-mCherry NE signal, indicating a reduced nuclear volume. Orange (nuclear) and blue (cytoplasmic) circles indicate areas used for quantification of the GFP signal. Nuclear projection areas were determined in the mCherry-channel using automated segmentation (see Materials and Methods) (scale bar: 5μm). B) Quantification of nuclear/cytoplasmic signal shows a significant leakage of NLS-GFP into the cytoplasm and thus indicates a shutdown of active nuclear transport already after 30 min of ED. The observed mean log2 fold change were: 3.276 (n=623) under control conditions; 0.662 (n=584) after 30 min; 0.3654 (n=722) 60 min and 0.2144 (n=604) >150 min after ED (all adjusted p-values <0.0001, ordinary one-way ANOVA and Tukey’s multiple comparison test). C) FRAP-recovery half-life times of nuclear signal from freely diffusing GFP at various time points after ED are significantly longer during ED as compared to control conditions (red dots) (p-value <0.0001, two-sided unpaired t-test) indicating a general down regulation of passive diffusion under these conditions. Passive nuclear diffusion of free GFP reaches a minimum after ~1 hour of ED and subsequently recovers slightly, pointing to cellular adaptation. The blue area shows the timepoint at which cryo-EM grids were prepared for structural analysis of ED NPCs. D) Histogram of quantified nuclear projection areas measured in segmented mCherry-channel of life-cell imaging as shown in (A) reveal a significant shift towards smaller values during ED (blue curves) as compared to control conditions (red curve) indicating a general loss of nuclear volume during ED that also manifest in NE wrinkling as seen in (A) (all adjusted p-values are <0.0001, one-way ordinary ANOVA and Holm-Sidak’s multiple comparison test with n=1056 control, n=1168 30min, n=2153 60min and n=1255 >150min after ED).

ED was shown to generally reduce cellular and nuclear volumes *S. pombe* cells (*28*, *29*, *31*). We therefore hypothesized that changes of diameter of the NPC could be a result of a reduced nuclear size that may reduce mechanical strain imposed onto the NPC scaffold by the nuclear membranes. As a proxy for nuclear size we quantified the median nuclear projection surface and indeed found a highly significant reduction from 71 μm^2^ to <60 μm^2^ under conditions of ED (**Fig. 4D**), while NE staining based on Nup60-mCherry fluorescence indicated NE wrinkling (**Fig. 4A**). This data therefore points to a general shrinkage of nuclei during ED. For a spherical nucleus, the observed changes in nuclear projection area correspond to a reduction of ~15% in nuclear surface area and ~25% in nuclear volume, which would be sufficient to cause a loss of NE tension (*50*) and may relax NPC scaffold into the constricted conformation (**Fig S12**).

## Discussion and Conclusion

Here we have investigated the compositional and conformational plasticity of NPC architecture in intact cells. We demonstrate that SpNPC scaffold exhibits an unexpected subcomplex arrangement that is breaking the long-standing dogma of a three ringed architecture. Similar to vertebrates and green algae, two concentric Y-complex rings form the NR. On the cytoplasmic side, eight individual cytoplasmic Y-complex vertices that do not exhibit any head-to-tail connection and thus do not form a ring. Although we cannot entirely exclude that the Y-complex tail is flexible at the cytoplasmic side and was thus not resolved during averaging, several lines of evidence argue against this. Previous biochemical analysis was suggestive of less tightly associated tail and vertex portions of the Y-complex in *S. pombe* (*41*). Another investigation suggested a non-isostoichiometric assembly of Y-complex members *in vivo* (*39*) and structural analysis of the Y-complex from yet another fungus, namely *Myceliophthora thermophila*, had demonstrated *in vitro* that Nup145C forms a stable fold and associates with the vertex in absence of Nup107 (*51*). Although the Y-complex does contain hinges (*32*, *51*–*54*) that are likely important to facilitate large scale conformational changes, the Nup189C-Nup107 interface is not known to be flexible. Taken together with the fact that the observed electron optical density sharply declines at the respective site, it is very likely that the interface between SpNup189C (HsNup96) and SpNup107, which was thought to be conserved, is not formed in the cytoplasm but only in the nucleoplasm. A recent study forced SpNup107 to the cytoplasmic side by expression of a SpNup189C-SpNup107 fusion protein which led to re-localization of SpNup131 to the cytoplasmic side (*39*), and thus further supports the here observed absence of the cytoplasmic Y-complex tail. How precisely *S. pombe* cells spatially segregate the two different types of Y-complexes remains uncertain. Our survey of public databases for splice variants, post-translational modifications and homologous structures did not yield significant clues. In contrast to vertebrates (*35*, *44*), *S. pombe* Ely5 is a member of both the nuclear and the cytoplasmic Y-complex vertex. It appears plausible that ELYS acquired additional functional domains during evolution of open mitosis in metazoans such as the C-terminal disordered region and AT-hook to tether Y-complexes to NPC seeding sites on chromatin and consequently to the nuclear side of the NE.

We further show how conformational changes in NPC architecture mediate its constriction and dilation within intact cells in response to a defined physiological cue, namely the energy status of the cell. ED leads to a massive constriction of the central channel that results in a ~2-fold loss in volume and is concomitant with a reduction of passive diffusion across the NE, while active nuclear transport is completely shut down. If the observed reductions of molecular exchange are directly or indirectly related to the NPC constriction remains challenging to address, given the manifold processes occurring in cells entering quiescence in response to ED (*28*–*30*, *55*, *56*). It however appears plausible that a reduction of the nuclear pore central channel volume limits the diffusion rate. In fact it has been suggested that NPCs reduce the diffusion rate of passively translocating molecules in response to their molecular size, rather than showing a strict size exclusion threshold (*57*). It has been further shown that active nuclear transport does not enhance passive diffusion (*58*, *59*) and several studies have shown that cytoplasmic diffusion of small proteins, such as soluble mCherry, is not significantly affected during ED (*28*). Finally, a recent study showed that the uptake and partitioning of both passively diffusing and nuclear transport factor (NTF)-like molecules by FG-domain *in vitro* is directly dependent on the their concentration (*60*). All of which agrees well with our findings. It therefore is plausible that a constricted central channel volume leads to an increased local FG-domain concentration which in turn limits the passive diffusion of molecules of a constant size, similar to the diffusion limitation observed in response to increasing molecular size under control conditions.

Peripheral channels are thought to be important for the nuclear import of inner nuclear membrane proteins (*11*–*13*). Here we observed around 3-4 nm wide lateral gaps between the individual spokes of actively transporting NPCs. Notably, our data processing workflow yields an average of conformation under the respective conditions and individual spokes are even more dynamic ((*6*) and this study). Therefore, it is plausible that the opening and closing of peripheral channels may regulate the translocation of inner nuclear membrane proteins.

Based on crystal structures of fragments of the Nsp1 (HsNup62) complex, it had been previously suggested that NPCs undergo dilation cycles that involve refolding and alternative configurations of the coiled-coil domains of the complex (*61*). The conformational changes observed in this study are very different. They do not necessitate a rearrangement of subcomplex folds but are rather based on large scale movements (**Videos S4-S7)**. Such movements may also be relevant during NPC assembly or turnover, where significant smaller diameters have been observed (*4*, *8*, *9*).

How ED mechanically leads to NPC constriction remains to be further addressed in the future. It appears likely that a reduced nuclear volume relieves NE tension and in turn allows NPCs to constrict (*50*). At this point, we cannot exclude additional factors such as the previously reported cellular pH-change during ED (*28*, *29*) or the shut-down of active nuclear transport itself to have an effect on NTF occupancy and NPC conformation. However, mechanical tension on the NE and active nuclear transport are certainly diminished during NE or NPC isolation. Therefore, previous structural analysis of such preparations has yielded structures that correspond to the most constricted conformation at the very end of the scale.

In conclusion we show that NPCs within livings cells populate a much larger conformational space and thereby confirm their importance as regulators of nucleocytoplasmic transport in response to environmental cues in living organisms on a cellular level. Hence our study highlights the power and importance of *in cellulo* structural analyzes to study such crucial physiological processes at the macromolecular level within the relevant cellular environment.

## Supplementary Material

Materials and Methods

Supplementary figures 1-12

Supplementary tables 1-3

Supplementary videos 1-7

## Author contributions

CEZ conceived the project, designed and performed experiments, acquired all types of data, designed and established data analysis procedures, analyzed all types of data, wrote the manuscript; MA conceived the project, designed and performed experiments, acquired data, analyzed data, wrote the manuscript; VR analyzed data, wrote the manuscript; SG designed and performed experiments, acquired data; AOK analyzed data; IZ designed and performed experiments; AH designed and performed experiments; JM designed experiments, supervised the project, JK conceived the project, designed and established data analysis procedures, analyzed data, supervised the project, wrote the manuscript; MB conceived the project, designed experiments, supervised the project, wrote the manuscript.

## Data availability

Associated with the manuscript are accession numbers EMD-11373, EMD-11374, EMD-11375. Integrative models will be deposited into the PDBDEV upon publication. The code, along the input files, for the modeling will be deposited in Zenodo upon publication.

## Acknowledgements

We thank Beata Turoňová, Nikola Kellner, Wim Hagen, Felix Weiss, Christian Tischer, Janina Baumbach, Ed Hurt, Edward Lemke as well as all the members of the Mahamid, Kosinski and Beck laboratories for advice and support. We thank Ed Hurt, Buzz Baum, Thomas Schwartz and Christian Häring for providing yeast strains. We acknowledge support from the Electron Microscopy Core Facility, the Advanced Light Microscopy Facility and IT services of EMBL Heidelberg. MA was funded by an EMBO a long-term fellowship (ALTF-1389–2016); JM received funding from the European Research Council (ERC 3DCellPhase^−^ 760067). MB acknowledges funding by EMBL, the Max Planck Society and the European Research Council (ComplexAssembly 724349).

