## Supplementary Material for "Nuclear pores constrict upon energy depletion"

### Supplementary Materials

#### Materials and Methods

##### *S. pombe* culture and cryo-EM grid preparation

Frozen stocks of *S. pombe* cells were freshly thawed and maintained on YES-agar plates (YES-broth from Formedium +20 g agarose/L) at 30 °C for maximum three days and re-streaked on fresh YES-plates prior to liquid culture inoculation. 5-10 ml YES-broth were inoculated with individual colonies and incubated over night at 30 °C shaking at 200 rpm, where $OD_{600}$ was kept below 1.0. Cryo-EM grid preparation was performed with a Leica EM GP plunger with a set chamber humidity of >90% and temperature of 30 °C. 3.5-4 µl cell suspension with previously adjusted $OD_{600}$ to 0.3 were applied to Cu200 or Au200 mesh R2/1 $SiO_2$ grids (Quantifoil) glow discharged on each side for 45 sec prior to sample application. The grids were blotted with Whatman 597 paper for 1-2 sec and plunge-frozen in liquid ethane or mix of ethane-propane (60% and 40%) at -189 °C or -194 °C, respectively. A list of all yeast strains and their source used in this study can be found in **Table S2**.

##### *C. thermophilum* culture and cryo-EM grid preparation

*Chaetomium thermophilum* (La Touche) *var. thermophilum* (from DSMZ, Braunschweig, Germany (No. 1495)) mycelium was maintained on CCM-plates as described in (*63*). For cryo-EM grid preparation Au200 mesh R2/1 $SiO_2$ grids (Quantifoil) were glow discharged on both sides for 45 sec and placed on the CCM-plates at the edge of the mycelium growth rim with the $SiO_2$ foil facing up. Plates were incubated at 55 °C until the mycelium covered at least half of the grid, typically 2-3.5 hours. Grids were then mounted in the chamber of a Leica EM GP plunger with a set chamber humidity and temperature at >90% and 55 °C respectively. Prior to plunge freezing in liquid ethane at -186°C, 3.5 µl plunging phosphate buffer (5% trehalose, 0.013 M $Na_2HPO_4$, 0.045 M $KH_2PO_4$, pH 6.5) was applied to the sample and blotted for 2 sec with Whatman 597 paper.

**Generation of knockout and fluorescently tagged *S. pombe* strains**

A *nup37Δ* knockout cassette containing a clonNat resistance marker corresponding to pFA6a-natMX6 (*64*), flanked by 70 basepair long homologous sequences to the nup37 gene was synthesized by Geneart and transformed in a K972 h- *wild type S. pombe* strain (kindly provided by the Häring lab at EMBL Heidelberg, Germany) after a modified protocol from (*65*). The *nup37Δ-ely5Δ* strain was generated by crossing the *nup37Δ* strain with a *ely5Δ* strain (*43*), kindly provided by the Schwartz lab at MIT Cambridge, MA, USA. Sporulation was triggered on sporulation plates prepared from SpoVB salts containing 8.2 g NaAc, 1.9 g KCl, 2.9 ml $MgSO_4$ 1 M, and 4.1 ml 5 M NaCl and 20 g agarose in 1 L $H_2O$. Tetrads of sporulated colonies were dissected in 40 µl sorbitol 1M with 10 µl zymolyase-T100. Selected colonies were replica plated on selection plates containing 100 µg/ml clonNat (Jena Bioscience) or 100 µg/ml G418 (Sigma-Aldrich). The double gene knockout was confirmed by PCR.

To generate the strain CZ001 expressing ubiquitous free super-folder GFP (sfGFP) with a Nup60-mCherry marker for FRAP-analysis, a Nup60-mCherry tagging cassette was amplified by PCR from isolated genomic DNA of the GD250 strain (*66*), kindly provided by the Baum lab at MRC LMCB London, UK. The product was transformed into the sfGFP expressing strain, AV0890 (*67*) from Yeast Genetic Resource Center (YGRC), as described above.

**Energy depletion of *S. pombe* cells**

Overnight cultures of *S. pombe* cells were grown at 30 °C in 5 ml dextrose-free Edinburgh minimal medium (EMM) supplemented with 20 mM glucose (glucose-control medium) while cell density was kept below $OD_{600}$ 0.8. Cells were collected and adjusted to obtain a cell amount of around 1 ml cells corresponding to $OD_{600}$ ~0.3 by centrifugation in Eppendorf tubes at 4000 rpm for 3 min at room temperature. Cells were then washed 3 times with 1 ml energy depletion buffer (20 mM 2-deoxy-glucose (Sigma-Aldrich) in dextrose free EMM and 10 µM antimycin A (Sigma-Aldrich)) and re-suspended in 1 ml energy depletion buffer. Control samples were washed 3 times in glucose control medium. For EM-grid preparation cells were incubated for 1 hour at 30 °C on a table-top shaker shaking at 900 rpm. The cell concentration was adjusted to an $OD_{600}$ of 0.2 and EM-grid preparation was performed as described above.

**Life fluorescent microscopy and FRAP imaging**

Ibidi µ-Slide 8-well Ibidi-treat dishes (Ibidi) were coated with 100 mg/ml Concanavalin A (Sigma-aldrich) in PBS for 10-40 min at room temperature and washed 3 times with $ddH_2O$. 300 µl of energy depleted and 15 µl of control cells were seeded into the Ibidi µ-Slide dish. Control samples were toped up with 150 µl glucose control medium to obtain a lower total amount of seeded cells to prevent cells from overgrowing during the following experiments. Time of energy depletion was measured from the first wash in energy depletion buffer, typically ~12 min before finalizing the final wash and seeding 8-well dishes.

Life cell imaging of CZ001 strain and FRAP experiments of GD250 strains were both performed on a laser scanning confocal microscope Zeiss LSM-780 equipped with incubation chamber maintained at 30 °C. All images were acquired with a Plan-Apochromat 63x/1.40 Oil DIC objective.

For life cell imaging of CZ001 1532x1532 pixel z-stacks were acquired with a pixel size of 0.0887 µm, an optical section of 0.8 µm and z-step of 0.38 µm using sequential line scanning and two line averaging rounds with 0.426 µs pixel-dwell time. GFP was excited with 488 nm line of the Argon laser and detected with GaAsP spectral detector in the range 500-550 nm. mCherry was excited with a DPSS laser at 561 nm and detected with GaAsP spectral detector in the range 568-620 nm. Per image stack 15-17 z-slices were acquired to cover the entire *S. pombe* cells. In total 3 independent biological replicates were acquired in 3 independent experiments with each containing ~5 individual z-stacks. For energy depleted cells, z-stacks were acquired at time intervals of 30-50 min, 55-70 min and 150-180 min after energy depletion (after first wash with energy depletion buffer) and control conditions were acquired more than 2 hours after sample preparation.

FRAP-measurements of bleached nuclear and cytoplasmic GFP signal in the CZ001 strains expressing ubiquitous sfGFP and Nup60-mCherry were performed to measure passive nuclear transport and cytoplasmic diffusion respectively.

Passive nuclear diffusion was measured by individual FRAP-experiments performed in the GFP channel consisting of a time series of 140 152x90 pixel (13.6 µm x 8.1 µm) frames acquired over 138.45 sec with an interval of 1 sec between different frames, a frame time of 0.17 sec and a pixel size of 0.09 µm. The 488 nm line of the Argon laser was used both for bleaching and for excitation during time lapse imaging. Bleaching was performed after acquiring 10 initial images. One bleaching iteration of 1.2 sec was performed in a circular area with a diameter of 33 pixel (2.97 µm), previously assigned manually based on the Nup60-mCherry NE outline to cover the nuclear area (**Fig. S11A**). Per individual biological replicate independent time series were acquired on individual cells over a period 30-210 min after energy depletion, i.e. after the first wash with energy depletion buffer. In total 48-55 FRAP-experiments per biological replicate were acquired consecutively on individual cells with an interval of around 3-5min between the individual measurements. Control measurements were performed after completion of data acquisition on energy depleted cells i.e. >210min after sample preparation.

To measure cytoplasmic diffusion, FRAP datasets were acquired in the GFP channel with image dimensions of 80x30 pixel (13.6 µm x 5.1 µm), a pixel size of 0.17 µm and 140 frames over a period of 2.11 sec with a time interval between frames equal to 0.03s. A rectangular area covering around half of the *S. pombe* cytoplasm and excluding the nucleus was bleached after acquiring 20 initial images (**Fig. S11C**). One bleaching iteration was performed for 20 ms with 100% power of the 488 nm laser line and additional 40% power of the 405nm laser line. In 3 independent biological replicates each, 8 individual FRAP-experiments at time points, 30 min (27-44 min), 60min (55-68 min) and >150min (155-168 min) after ED were recorded on individual cells. Control measurements were performed after 180 min of sample preparation.

**Quantification of fluorescent microscopy images and FRAP-analysis**

To quantify the localization of NLS-GFP in GD250 cells, z-stacks were maximum projected and the ratio of the average pixel intensity of a circular area within the nucleus compared to a similar area in the cytoplasm were measured manually in Fiji (*68*) (**Fig. 4A**). The nuclear position was determined in the Nup60-mCherry channel. Cells that were not well attached to the 8-well dish and moving during z-stack acquisition or not entirely contained within the z-stack were removed prior to analysis and cytoplasmic areas of clearly visible vacuoles were avoided.

Nuclear projection areas were quantified by segmenting nuclei in the Nup60-mCherry channel in maximum projected z-stacks with excluded areas of cells moving or not entirely contained within the imaged volume by Ilastik (*69*) using the auto-context workflow. The areas of the segmented images were then quantified using Mathwork's MATLAB.
FRAP-curves were analysed with FRAPAnalzer 2.1.0 (*70*). In brief a double normalization was carried out normalizing against the background and the entire cellular surface area as reference. Recovery half-lifetimes were extracted by fitting a one exponential recovery equation to the normalized data.

**Cryo-FIB milling**
Plunge-frozen sample grids were FIB-milled in an Aquilos FIB-SEM (Thermo Fischer) as previously described (*4*). In brief, samples were sputter coated with inorganic platinum at 10 kV voltage and 10 mPa argon gas pressure for ~10 sec (Pt-sputtering). Subsequently a protective layer of organometallic platinum was deposited using the gas injection system (GIS-coating) for ~12 sec. If needed a second round of Pt-sputtering was performed to avoid charging during FIB-milling. Between 5 and 10 lamellae per grid were milled in a step-wise fashion and polished to a final thickness of <280 nm using decreasing FIB current steps of 1 nA, 0.5 nA 0.3 nA and 50 pA. Before unloading the sample, an additional layer of Pt-sputtering was deposited for 1 sec to avoid charging effects during subsequent imaging in the transmission electron microscope (TEM).

**Automated Tomogram acquisition**
The majority of *wild type S. pombe* tomograms (136/178) as well as all tomograms from knock-out strains were acquired on a Titan Krios G4 (Thermo Fischer), operating at 300 keV and equipped with a Gatan K2 Summit direct electron detector and energy-filter. All tilt-series (TS) were acquired in dose-fractionation mode at 4k x 4k resolution at a nominal pixel size of 3.45 Å. Automated TS acquisition was performed as described previously (*4*) using a dose-symmetric acquisition scheme (*71*) with an effective tilt range of -50° to 50°, considering a tilt-offset to compensate for the lamella angle (typically +/- 8-13° ) (*4*), a 3° tilt increment, a total dose of 120-150 e/$Å^2$ and target defocus of -1.5 µm to -4.5 µm.

The remaining 42 *wild type* tomograms and 15 of total 76 tomograms from energy depleted *S. pombe* cells were acquired on a Titan Krios (Thermo Fisher) equipped with a Gatan K2 Summit direct electron detector, energy-filter and a Volta potential phase plate (VPP) (*72*). Automated dose-symmetric tilt series acquisition was performed in 4k x 4k dose-fractionation mode with a nominal pixel size of 3.37 Å using a total dose of 100-125 e/Å$^2$ distributed over a tilt range of -60° to +60° or -50° to 50° with and a tilt step interval of 3° or 2° respectively, a defocus range of -2 µm to -4 µm and conditioning the VPP by exposure to the electron beam for up to 5 min. Additional 11 tomograms of ED *S. pombe* cells were acquired using the same parameters but without usage of the VPP.

The remaining 50 tomograms of ED *S. pombe* cells were acquired on a Titan Krios G4 (Thermo Fischer) operating at 300 keV equipped with a Gatan K3 direct electron detector and energy-filter operated in dose-fractionation mode and 5.7k x 4k resolution with a nominal pixel size of 3.425 Å. Automated dose-symmetric tilt series acquisition was performed with a nominal defocus range of -2 µm to -3.5 µm, an effective tilt range of -50° to 50° including a tilt offset to compensate for the lamella pre-tilt, a tilt step increment of 3° and a total dose of 133 e/Å$^2$. *C. thermophilum* tomograms were acquired as described above for *wild type S. pombe* tomograms with a total dose of 130-140 e/Å$^2$ and a target defocus range of -2 µm to -4 µm.

**Image pre-processing and tomogram reconstruction**

Images were pre-processed and CTF was estimated as described previously (*4*). Tilt series were aligned automatically in a tailored workflow using the IMOD package including the patch-tracking functionalities (*73*). In detail, 4-times binned (pixel size: 1.348 nm) and dose-filtered TS were low-pass filtered with a set of 7 empirically pre-defined low-pass filters and 7 initial patch-tracking attempts using 7x7 patches per tilt were performed in IMOD. The filter-set and patch tracking output yielding the lowest alignment residuals during an initial round of patch-tracking was selected for further iterative refinement. At each iteration the contour with the largest residual error was removed prior to a next round of patch-tracking. This process was repeated until the overall residual dropped below 0.5 pixel or until a ratio of known/unknown in the IMOD tilt align program dropped below a threshold of 10. The final aligned TS were reconstructed using the SIRT-like filtering option in IMOD (*74*) and manually inspected. Only tomograms yielding a high quality alignment, both, visually and in terms of overall residual error (typically smaller than 1 pixel) were used for 3D CTF correction using the phase-flipping option in novaCTF (*75*) and subsequent subtomogram averaging workflow.

**Particle identification and subtomogram averaging**

NPC coordinates and initial orientations were determined manually in 4-times binned SIRT-like filtered (*74*) tomograms as described earlier (*4*). Additionally, a model describing the lamella slab geometry was generated manually by picking the coordinates of all 8 corners of the lamella slab within each tomogram. Particle extraction, subtomogram alignment and averaging (STA) was performed with novaSTA (*76*) from 3D CTF corrected tomograms as described previously (*4*). Initial alignment of NPCs was carried out with in 8- and 4-times binned particles. After obtaining an initial 4-times binned average coordinates of the 8 individual spokes per NPC were assigned in the original tomograms for subunit sub-boxing. Only subunits with coordinates retained in the lamella slab geometry model were extracted to avoid the inclusion of subunits laying outside the FIB-milled lamella. At this step all VPP data was excluded from further STA for the *wild type* dataset. After an initial subunit alignment with 4-times binned particles, each subtomogram and its assigned orientation was inspected manually and misaligned or remaining false positive particles, i.e. subunits extracted outside of the FIB-milled lamellae, were removed. Subtomogram alignment was further continued using 4- and 2-times binned subtomograms and localized masks for the individual rings: cytoplasmic side, IR and NR. If needed the particle boxes were recentered around the individual rings to achieve better subtomogram alignment. In case of the *wild type* subtomogram average, particles with a low final cross correlation scores were removed before creating the final average. All subtomogram averages of *wild type* and knock-out strains were b-factor sharpened empirically (*35*) and filtered to the given resolution determined on gold-standard FSC calculations at the 0.143 criterion. For ED averages, instead of b-factor sharpening, the amplitude of the finial averages were matched to the amplitudes obtained from averaging the final aligned defocus data only (excluding all VPP data) with the EMAN2 (*77*) e2proc3d.py -matchto function, to overcome a previously described low-pass filter-like artifact occurring during subtomogram averaging of VPP data (*78*). Details about tomogram and particle numbers of each average can be found in **Table S3**.

**Difference map calculation**

To calculate the difference map of the *nup37Δ* and *nup37Δ-ely5Δ* knock-out to the *wild type* maps, knock-out whole pore assemblies were generated based on the *wild type* subunit coordinates and represented on the *wild type* membrane as follows: First the individual ring segments were filtered according to their resolution for *nup37Δ* cytoplasmic side: 27 Å, IR: 27 Å and NR: 31 Å and cytoplasmic side: 35 Å, IR: 35 Å and NR: 34 Å of *nup37Δ-ely5Δ* respectively (**Table S3 and Fig. S7**). The Individual maps were then fitted to copies of the corresponding *wild type* segments, which were previously filtered and scaled to match the respective resolutions and density distribution of the knock-out maps. The fitted segments were assembled to a complete subunit spoke and an 8-fold rotational symmetric assembly was generated based on the *wild type* coordinates. The matched and filtered *wild type* segments were assembled to 8-fold rotational symmetric pores based on the previously determined coordinates and the knock-out 8-fold assembly was subtracted from the *wild type* density to obtain a difference map in UCSF Chimera (*79*). For final representation the 8-fold knock-out protein assemblies were represented on the *wild type* membrane and overlaid with the corresponding difference map.

**NPC diameter and volume measurements**

NPC diameters were measured with an in-house MATLAB script based on the final coordinates and orientations obtained for each individual subunit during STA. First a feature of interest in the subunit average was identified in UCSF Chimera (*79*) by placing a 1-pixel sphere mask on the point of interest. The offset between the center of the average and the mask was then used to calculate the coordinates of the area of interest within each averaged subunit in respect to the original tomograms. To calculate the diameter of each individual NPC, only NPCs with a subunit-occupancy of five or more were considered. For each individual NPC vectors were derived connecting the opposing subunits. Based on those vectors the center of each NPC was defined as the center of the intersection area between all vectors corresponding to a given NPC. The average distance between the newly identified center and each respective SU was chosen as a representative NPC radius for the given pore. By this means, an accurate average NPC radius measurement for a specific feature of interest within each individual NPC is obtained. Statistical tests to compare NPC diameter differences across different conditions were performed using two-sided unpaired t-tests or Mann-Whitney test in Prism Graphpad 8. Central channel volumes were estimated by calculating the volume of two cones using the formula $V=1/3\pi(r_1^2r_1r_2+r_2^2)h1 + 1/3\pi(r_2^2+r_2r_3+r_3^2)h_2$ with average radii r obtained above and heights h measured from the final STA cryo-EM maps as indicated in **Fig. 8C**.

#### Homology modeling

Modeling templates were detected and selected using the HHpred server (*80*). Sequence alignments for modeling were refined in Swiss-PdbViewer (*81*). The models were built based on the templates and alignments using Modeller (*82*). The templates used for modeling are listed in **Table S4**.

#### Systematic fitting of SpNPC rigid bodies to cryo-EM maps

An unbiased global fitting approach was implemented utilizing several SpNPC structural models. Two of them were experimental X-ray structures previously published (*42*, *43*) while the remaining were homology modeled based on *S. cerevisiae* and *C. thermophilum* NPC components (*15*, *83–86*) (**Table S4**). All the aforementioned high-resolution structures were filtered to 40 Å prior the fitting. The resulting simulated model maps were subsequently fitted into individual segments of the SpNPC cryo-EM maps from this study by global fitting as implemented in UCSF Chimera (*79*). More precisely, structures of the Y-complex were fitted into cytoplasmic and NR segments of the cryo-EM SpNPC maps while IR model was fitted into IR. The individual segment maps used for fitting did not include any nuclear envelope density in order to eliminate the possibility of fits significantly overlapping with the membrane.

All fitting runs were performed using 100,000 random initial placements with the requirement of at least 30% of the simulated model map to be covered by the SpNPC density envelope defined at low threshold. For each fitted model, this procedure resulted in approximately 250-17,200 fits with non-redundant conformations upon clustering. The cross-correlation about the mean (cam score, equivalent to Pearson correlation) score from UCSF Chimera (*79*) was used as fitting metric for each atomic structure, similarly to our previous published works (*4*, *35*, *36*, *38*). The statistical significance of every fitted model was evaluated as a p-value derived from the cam scores. The calculation of p-values was performed by first transforming the cross-correlation scores to z-scores (Fisher's z-transform) and centering, from which subsequently two-sided p-values were computed using standard deviation derived from an empirical null distribution (based on all obtained non-redundant fits and fitted using fdrtool (*87*) R-package). Finally, the p-values were corrected for multiple testing with Benjamini-Hochberg (*88*). Figures were produced by UCSF Chimera (*79*) and UCSF ChimeraX (*89*).

**Integrative modeling**

Structural models of the SpNPCs were built using an integrative procedure (**Fig S4**) similar to the one we used previously for human and *S. cerevisiae* NPCs (*4*, *36*). First, a model of the dilated (actively transporting) NPC was constructed. The cytoplasmic Y-complexes and NR were built in a two-step procedure comprising "multiple fitting" and "refinement" steps. The homology models were divided into smaller rigid bodies (with cut points corresponding to boundaries of published crystal structures) and simultaneously fitted to the cryo-EM map using a custom methodology (*4*, *36*) implemented based on Integrative Modeling Platform (IMP) (*90*) version 2.13 and Python Modeling Interface (PMI) (*91*). In the multiple fitting step, first libraries of alternative non-redundant fits for each rigid body were generated based on the systematic fits described above. Then, a simulated annealing Monte Carlo optimization was run to generate configurations of all the rigid bodies by recombining the aforementioned fits using simulated annealing Monte Carlo method. To cover a large landscape of possible models, the optimization was run independently 10,000 and 40,000 times for the cytoplasmic side and NR respectively, with each run leading to a candidate model. To achieve convergence and sampling exhaustiveness (assessed by checking whether new better scoring models appear with extra steps), each run comprised by 130,000 (cytoplasmic side) and 271,000 (NR) Monte Carlo steps at decreasing temperatures. Higher number of steps and models for NR were used because of the higher number of rigid bodies in this ring. The scoring function for the optimization was a linear combination of the EM fit restraint represented as the p-values of the pre-calculated domain fits (from systematic fitting as described above), clash score (SoftSpherePairScore of IMP), connectivity distance between domains neighboring in sequence, a term preventing overlap of the protein mass with the nuclear envelope, a restraint promoting the membrane binding loops of Nup131, Nup132 and Nup120 to interact with the envelope, implemented using MapDistanceTransform of IMP (predicted by similarity to known or predicted ALPS motifs in human and *S. cerevisiae* homologs (*1*, *92*) and the restraint for Ely5 localization in the region identified based on the difference EM maps to the *nup37Δ* and *nup37Δ-ely5Δ* NPC maps. During the optimization, the structures were simultaneously represented at two resolutions: in Cα-only representation and a coarse-grained representation, in which 10-residue stretches were converted to a bead. The 10-residue bead representation was used for all restraints to increase computational efficiency except for the domain connectivity restraints, for which the Cα-only representation was used. Since the p-values used for the EM restraint were derived from the original EM full atom fitted models generated with UCSF Chimera, the EM restraint can be regarded as an EM restraint derived from the full atom representation. The model of the cytoplasmic side was built by using all Y-complex components including Ely5. Due to the higher complexity of NR, it was first built without Ely5, which was then added separately using the refinement protocol described below. For the NR, a restraint promoting Nup37 localization in the difference density to the *nup37Δ* NPC was also used.

In the refinement step, the top model of the cytoplasmic side and the top 10 models of NR were subjected to a Monte Carlo simulated annealing optimization of the rigid body rotations and translations but this time without using the libraries of alternative non-redundant fits, allowing the rigid bodies to move in the EM map with small rotation and translation increments. The scoring function consisted of a cross-correlation to the EM map (FitRestraint of IMP), domain connectivity restraint, the clash score, and the envelope exclusion and binding restraints as above.

Owing to high structural conservation of the IR, this ring was built without the "de novo" mode but instead by fitting a homology model of the entire IR built based on *S. cerevisiae* NPC, followed by the refinement with 50,000 Monte Carlo steps at a single temperature. The procedure was similar as for the cytoplasmic side and NR, including a restraint for membrane-binding loop of Nup155 (*36*) and a restraint enforcing that Ely5 interacts with Nup120 based on (*43*).

To generate the final models of the dilated and ED NPCs, the top-scoring models for CR, NR and IR for the dilated NPC where merged together and subjected to a multi-state refinement procedure, which optimized the NPC structure simultaneously in all three, dilated and ED, EM maps. This was necessary to prevent over-fitting to a particular map, which could generate artificial conformational differences between the resulting models, stemming solely from random variations in the EM density. The multi-state refinement procedure involved three parallel simulated annealing Monte Carlo replicas with each replica subjected to the restraints coming from a different EM map. The scoring function composed of the restraints as for the refinement of the individual rings except the *nup37Δ* and *nup37Δ-ely5Δ* NPCs difference map restraints. In addition, and uniquely to the multi-state modeling, a restraint that keeps the three models as similar to each other as allowed by other restraints was imposed under a parsimonious assumption that the models should be locally similar unless the cross-correlation with the EM map strongly indicates otherwise. This restraint was implemented as a network of elastic (harmonic) distance restraints connecting beads within 15 Å threshold to each other. Finally, another elastic network restraint was used to keep the Ely5-Nup120 orientation similar in all rings. The optimization was run 100 times (each with 210,000 steps at six temperature levels) and the top-scoring model out of the resulting 100 models was used as a representative model for the figures.

### Supplementary Figures

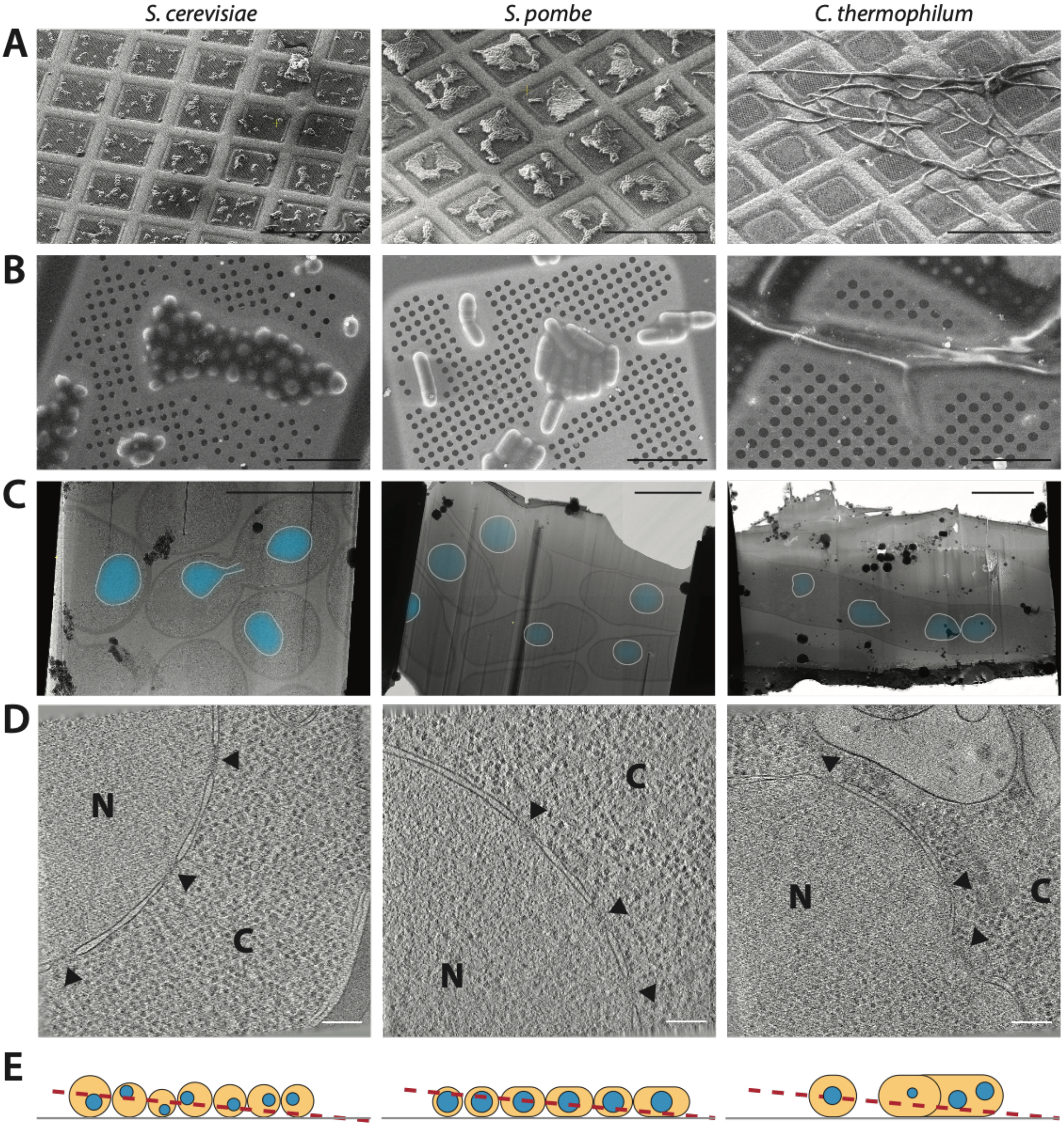


**Supplementary figure 1. FIB-milling of different fungi.** A) Grid overview by scanning electron microscopy (SEM). While *S. pombe* and *S. cerevisiae* cells form assemblies of several cells, *C. thermophilum* mycelia are more separated (scale bar 200 µm). B) Higher magnification scanning electron micrographs of an individual yeast cell assemblies, and an individual *C. thermophilum* filament (scale bar 20 µm). C) Transmission electron micrographs of FIB-milled lamellae overlaid with the segmented nucleus (blue) and NE (white). *S. cerevisiae* cells have a relatively small nucleus with respect to cell diameter (see also D) and as a consequence not all *S. cerevisiae* cells are milled at a z-height that includes the nucleus within the lamella-slab, in contrast to *S. pombe* cells. Mycelia of *C. thermophilum* contain several nuclei with often irregular shape (scale bar 5 µm). D) Sections through reconstructed and filtered tomograms. NPCs are indicated by black arrow heads, N marks the nucleoplasm and C marks the cytoplasm. *S. pombe* tomograms are contrastic with an average of ~3.9 NPCs per tomogram whereas in *S. cerevisiae* only an average of ~2.1 NPC per tomogram are found (scale bar 150nm). E) Cartoon indicating the different cellular geometry of fungal cells and the consequences for FIB-milling. The rod-shaped geometry of *S. pombe* cells is ideal for FIB-milling, especially to target high number of nuclei. The filament geometry of *C. thermophlium* imposes a considerable challenge to produce stable lamellae. All images containing *S. cerevisiae* data were generated based on (*4*).

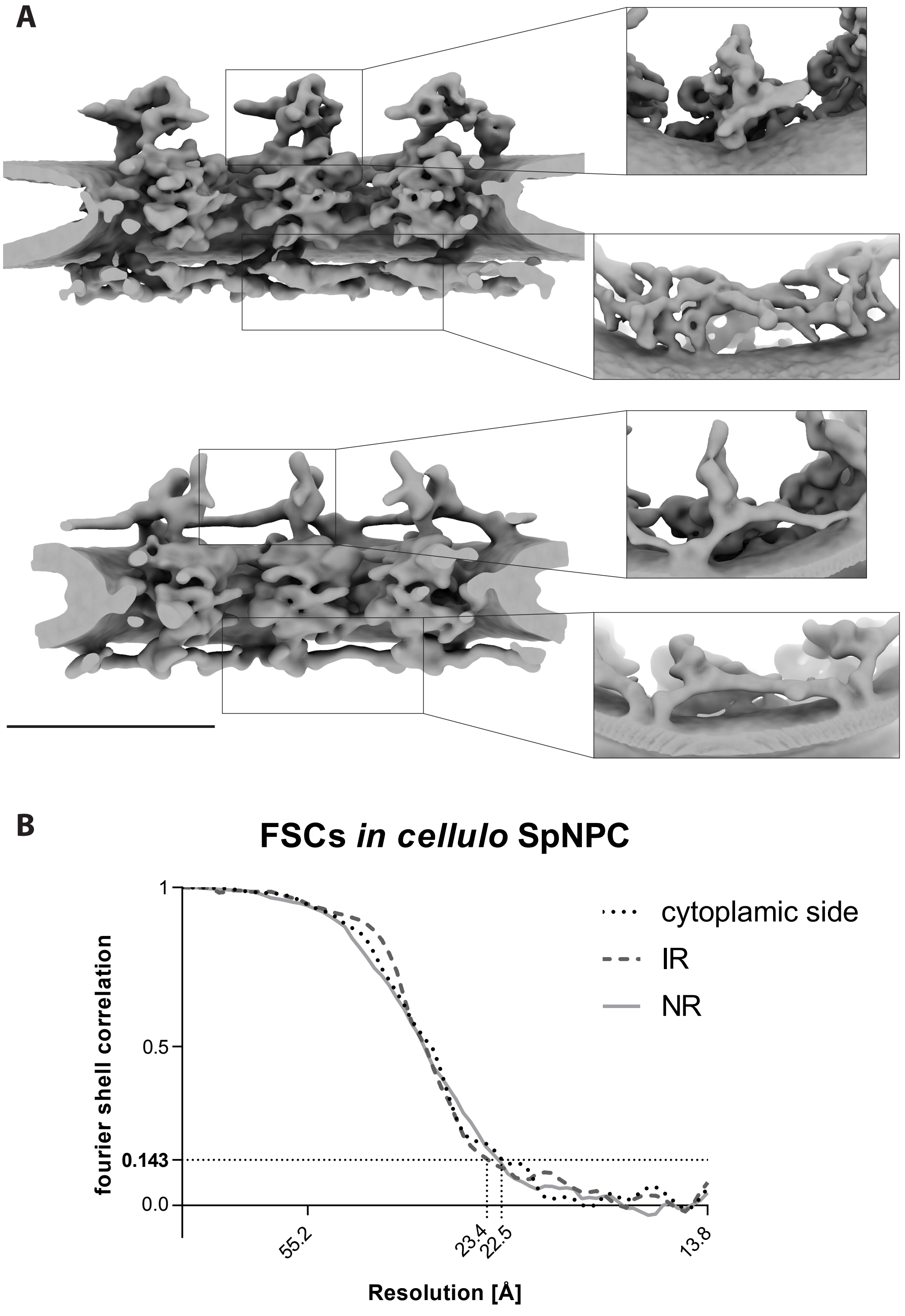


**Supplementary figure 2. Detailed features of the *in cellulo* cryo-EM map of yeast NPCs.** A) Comparison of the *in cellulo S. pombe* and *S. cerevisiae* cryo-EM maps. The *S. pombe* map is better resolved and reveals additional features on the protein domain level (scale bar: 100nm). B) Fourier shell correlation (FSC) curves of the SpNPC cryo-EM map reveal an overall resolution of better than 23.4 Å using the 0.143 criterion for the cytoplasmic side, inner (IR) and nuclear ring (NR) that were refined separately on the asymmetric unit level.

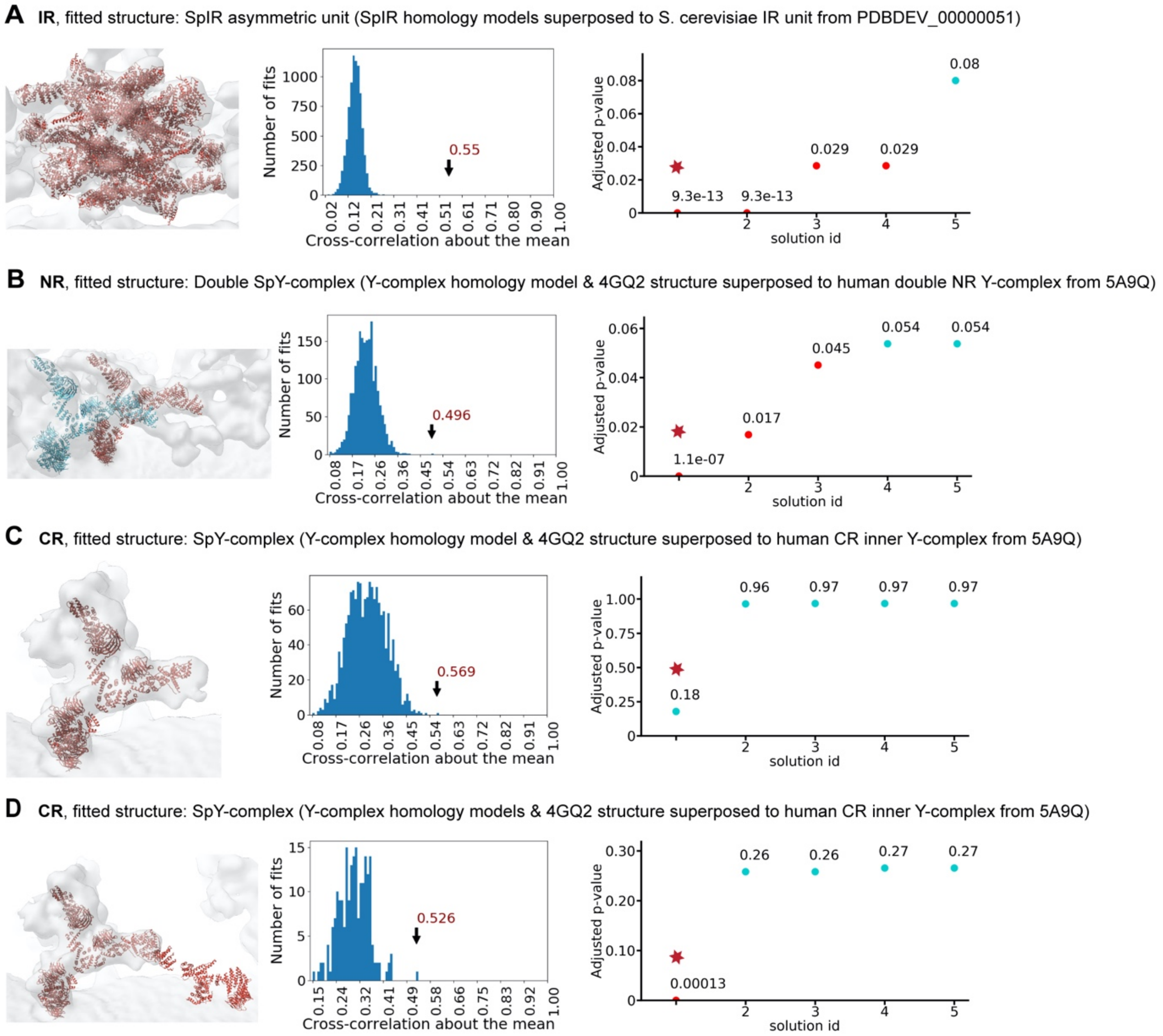


**Supplementary figure 3. Systematic fitting of high-resolution structures and homology models into the cryo-EM map of the actively transporting SpNPC.** Each panel shows the visualization of the top fit (left), the histogram of the cross-correlation about the mean (cam) scores (middle), and a plot of the top five p-values (right). In the p-value plots, the statistically significant fits are colored red (p-value < 0.05; p-values were calculated using the two-sided test as implemented in fdrtool R-package, as described in Materials and Methods). The top fits are indicated in the histograms with an arrow and the score value and in the p-value plots the top fits are indicated with a red star. The number of sampled fits used to calculate p-values after clustering of similar solutions was 8371 (A), 2030 (B), 1605 (C) and 210 (D). A) Fitting of the IR. All asymmetric unit homology models were superposed to the integrative model of the single spoke of the IR unit from PDB-DEV_ID 00000051 and the resulting composite model was used for the fitting. The two best scoring fits represent the orientations of the C2 symmetric rigid body rotated by 180°. B-D) Fitting of the cytoplasmic side and NR.

The crystal structure of Nup120-Nup37 (PDB-4GQ2) and the SpY-complex homology models (covering Nup120-Nup189c-Sec13-Seh1-Nup85-Nup107, Nup107-Nup131/Nup132 and Nup131/Nup132 β-propeller) were superposed on the human Y-complex models (PDB-5A9Q) and systematically fitted. The two composite models of NR SpY-complexes shown in (B) were fitted as a single rigid body and colored differently for visualization purposes only (outer Y-complex blue, inner Y-complex red). All components were fitted to the cryo-EM maps of the individual cytoplasmic side, IR and NR segments.

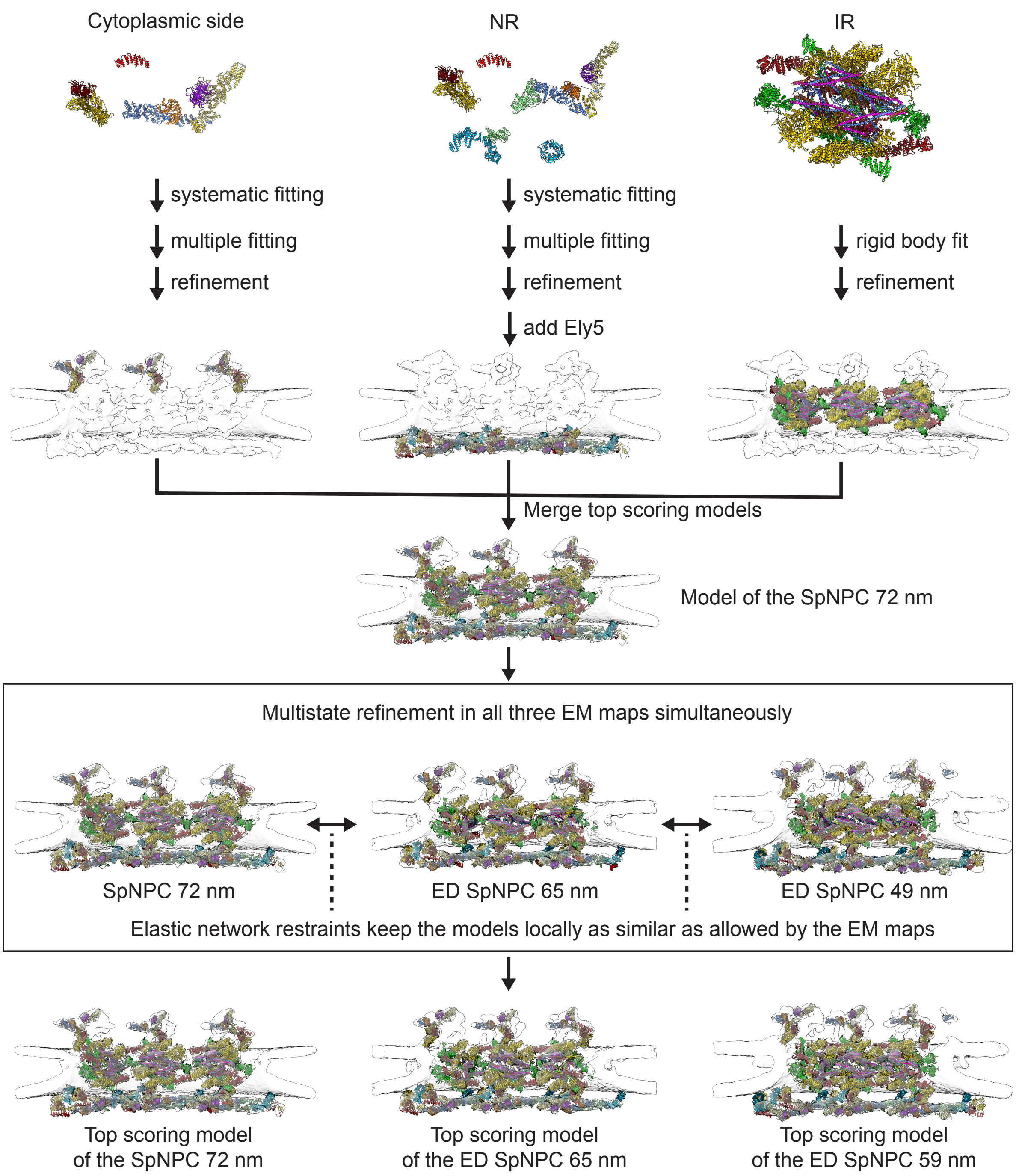


**Supplementary Figure 4. Integrative modeling procedure used to build the models of the SpNPC and ED SpNPCs.** First, for the cytoplasmic side and NR, individual structures were fit independently to the EM map leading to large libraries of alternative fits (*systematic fitting*). The resulting alternative fits were then used to build models comprising all structures (*multiple fitting*). The *refinement* step further optimized the fits using a more fine-grained optimization protocol. For NR, Ely5 was added after the refinement of the remaining structures through another round of refinement. The *rigid body fit* of the IR indicates that for this ring, the starting point was a homology model of the IR built based on the *S. cerevisiae*

NPC. Further details of the procedure are described in Materials and Methods. The cryo-EM maps are shown as transparent surface. The structural models are shown in cartoon representation and colored as in **Fig. 3**. The quality of the final fits in close-up views is depicted in the Supplementary Video 1. Abbreviations: inner ring (IR), nuclear ring (NR), ED SpNPC – Energy Depleted SpNPC.

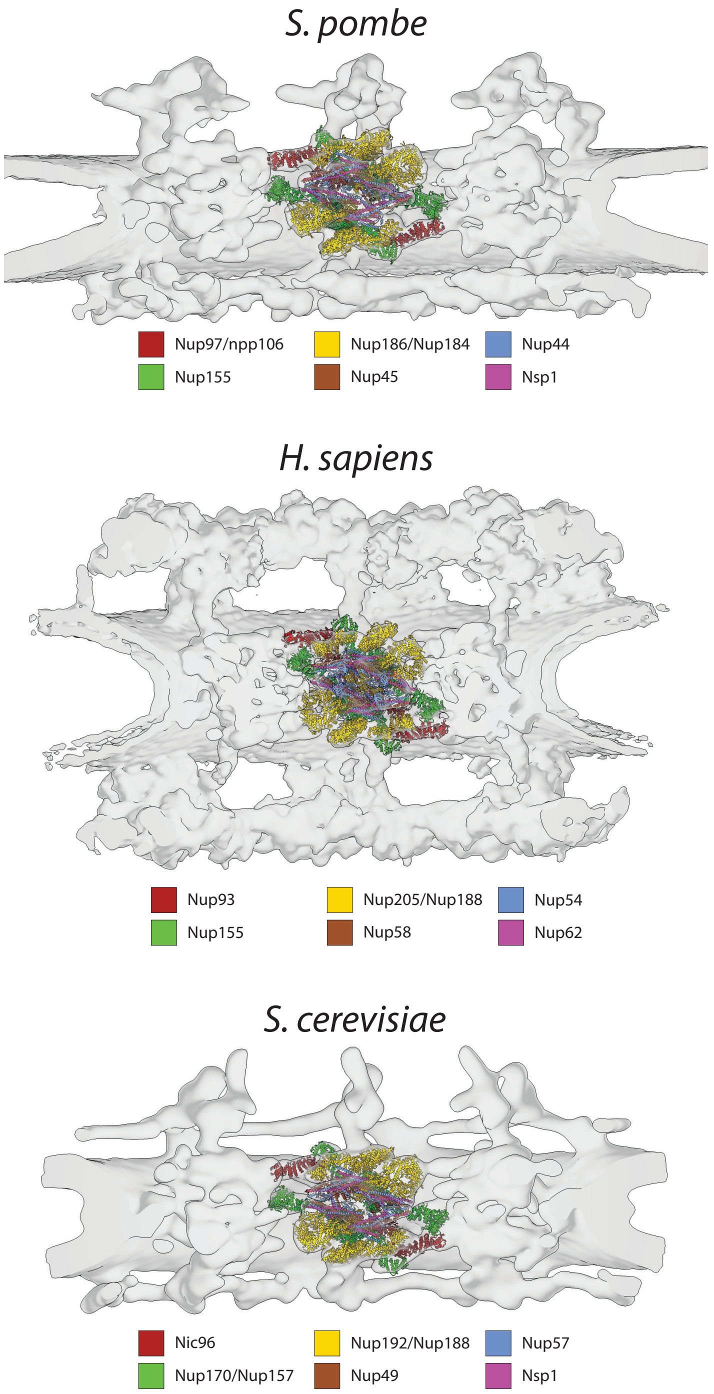


**Supplementary figure 5. The inner ring scaffold architecture is conserved from yeast to humans.** Comparison of the *S. pombe* IR model fitted to the cryo-EM (top) with the human model PDB-5IJO fitted in EMDB-3103 (middle) and *S. cerevisiae* IR model PDB-DEV_ID 00000051 fitted to EMDB-10198 (bottom). All three IRs follow the same architectural blueprint with four copies of the SpNup97/HsNup93/ScNic96 subcomplex (red, green and yellow) serving as anchoring platform for the four copies of the triple coiled-coil channel nucleoporin heterotrimer (CNT) (also called Nsp1/HsNup62-subcomplex) (brown, blue and purple) pointing towards the central channel. The structural models explain the vast majority of the observed electron optical densities.

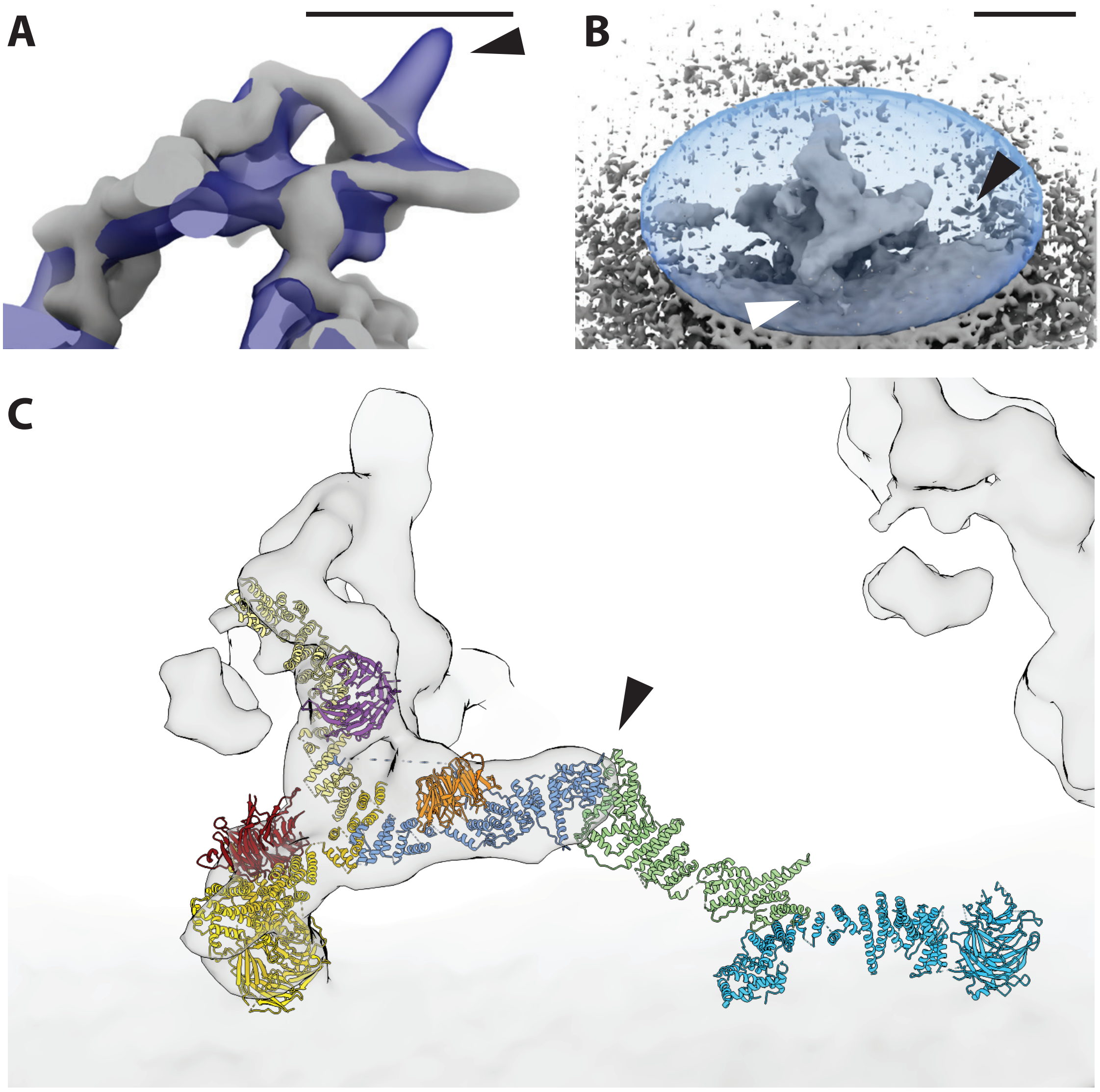


**Supplementary figure 6. The cytoplasmic side of the SpNPC contains of a *bona fide* Y-complex vertex.** A) The overlay of the *S. pombe* (grey) and *S. cerevisiae* (blue) mRNA export platform reveals an overall similar architecture despite the absence of the dynein arm (arrowhead; scale bar 20 nm). B) Refinement of the cytoplasmic protein entity with a large ellipsoidal mask did not recover any additional electron optical density accounting for either the Y-complex tail (black arrowhead) or the membrane binding β-propeller of a preceding Y-complex tail (white arrowhead), thus arguing against a head-to-tail arrangement (scale bar 20 nm). C) Fitting of the nuclear Y-complex arrangement to the cytoplasmic side identifies a significant fit of the entire Y-complex (see also **Fig. S3D**). The tail is thereby positioned outside of the observed density that sharply declines at the interface between Nup189C and Nup107 (arrowhead, same color code as in **Fig. 2**).

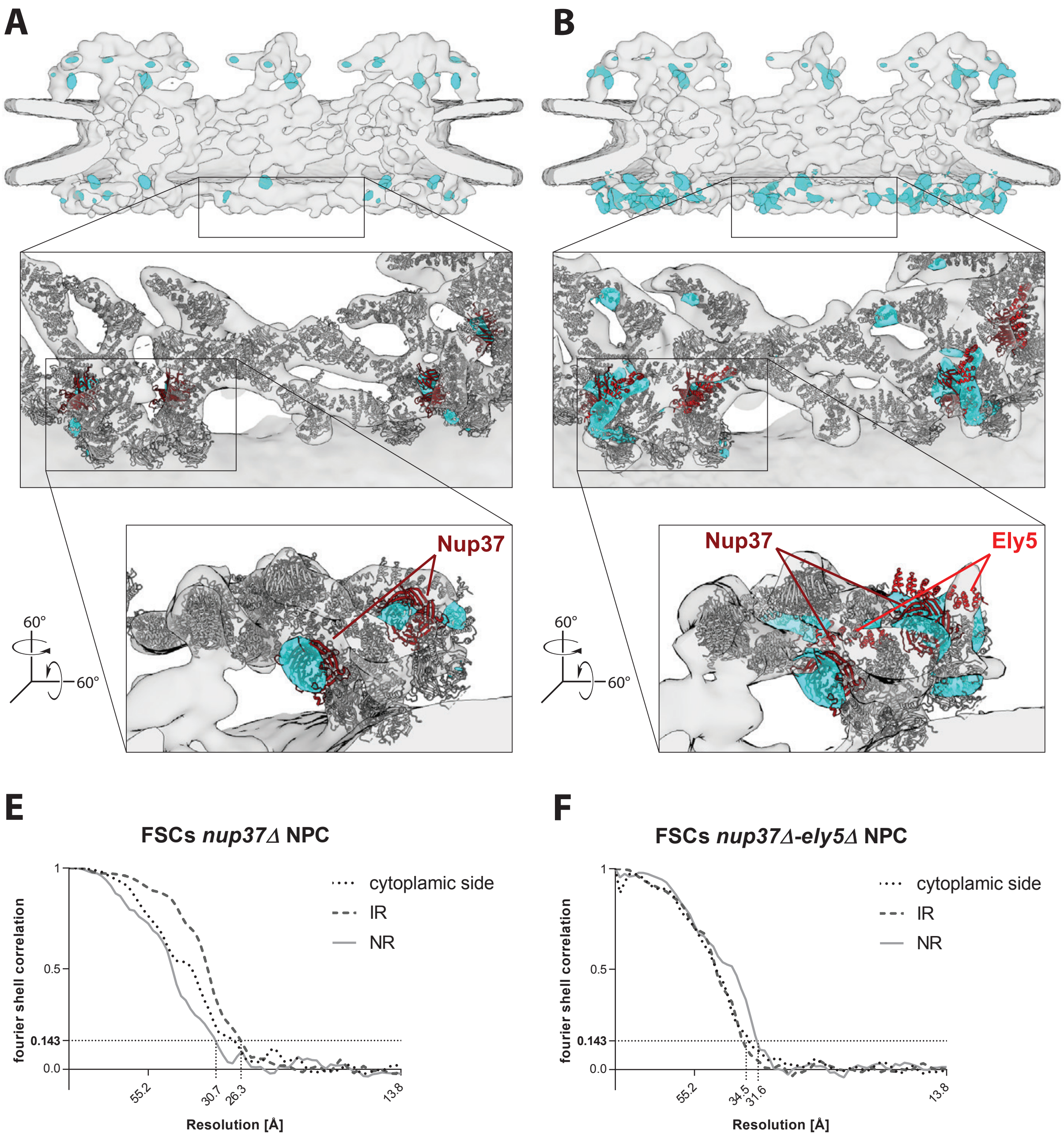


**Supplementary figure 7. *nup37Δ* and *nup37-Δely5Δ* cryo-EM maps confirm positioning of the Y-complexes and identify Ely5 as cytoplasmic Nup.** A) cryo-EM map of the *nup37Δ* SpNPC (light grey) overlaid with a difference map (cyan) of the *wild type* and *nup37Δ* maps (see Materials and Methods) and the composite model of the Y-complexes (grey). No differences are apparent in the IR. At the NR, density is missing at the expected location of Nup37 (red). Very few rather small differences are observed in addition, e.g. in the Nup85 arm, hinting to a potentially altered conformation or increased flexibility. B) Same representation as in A) for the *nup37Δ-ely5Δ* double knockout map. On the nuclear side, pronounced differences at the expected location of Nup37 and Ely5 (red) are identified. Additional differences in the region of the Y-complex vertices point to an increased flexibly and slight conformational change of this structure under *nup37Δ-ely5Δ* knockout conditions. E) FSC curves of the individual *nup37Δ* cryo-EM maps. F) FSC curves of *nup37Δ-ely5Δ* cryo-EM maps.

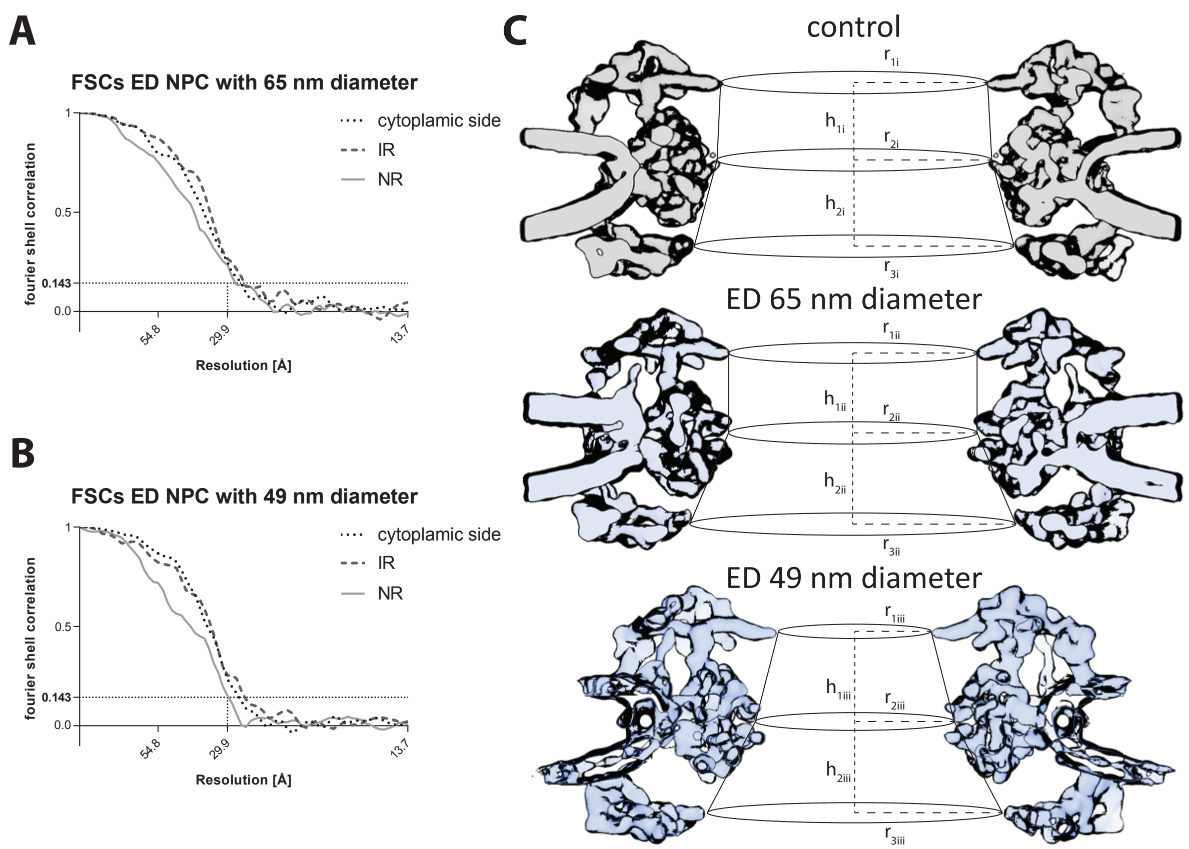


**Supplementary figure 8. FSC curves and estimation of central channel volume.** FSC curves of the classes with diameters 65 nm (A) and 49 nm (B). C) The volume of the central channel of actively transporting (control), and energy depleted NPCs with intermediate and constricted diameter (top to bottom) was estimated from the subtomogram averages by calculating the volume of two truncated cones with radii $r_1$, $r_2$ and $r_2$, $r_3$ respectively and the corresponding heights ($h_1$, $h_2$). $r_1$ corresponds to the cytoplasmic diameter at the likely anchor point of the cytoplasmic FG-repeat domains, $r_2$ represents the diameter between the potential FG-anchor points of the inner ring subcomplexes and $r_3$ describes the NR diameters at the inner Nup85-arm tip lining the central channel. For simplicity, this model neglects the opening of the peripheral channels within the IR upon dilation and will thus slightly underestimate the volume increase. The following dimensions were measured, control: $r_{1i}$=33.6 3nm, $r_{2i}$= 35.76 nm, $r_{3i}$=40.11 nm, $h_{1i}$=17.94 nm and $h_{2i}$=18.63 nm resulting in central channel volume $V_i$=152,317.86 nm$^3$. ED 65 nm diameter: $r_{1ii}$=29.65 nm, $r_{2ii}$= 32.37 nm, $r_{3ii}$=38.86 nm, $h_{1ii}$=17.81 nm, $h_{2ii}$=18.49 nm and $V_{ii}$=127,723.25 nm$^3$. ED 49 nm diameter: $r_{1iii}$=21.56 nm, $r_{2iii}$= 24.5 nm, $r_{3iii}$=35.3 nm, $h_{1iii}$=19.18 nm, $h_{2iii}$=19.18 nm and $V_{iii}$=86,456.83 nm$^3$.

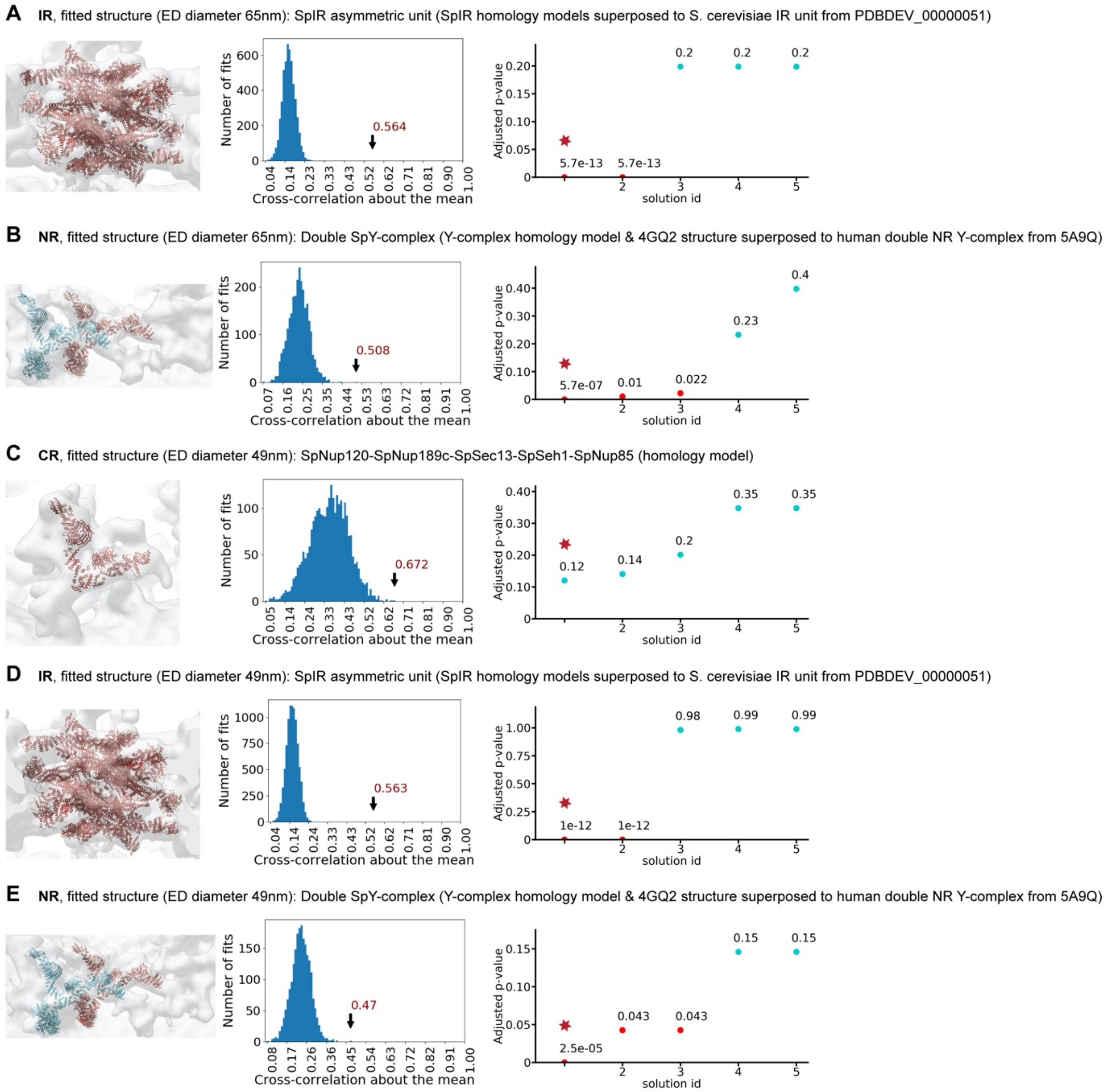


**Supplementary figure 9. Systematic fitting of inner and outer ring components into the energy depleted SpNPC maps (diameters 65 nm & 49 nm).** Same as **Fig. S3** but for the two classes of constricted NPCs. The number of sampled fits used to calculate p-values after clustering of similar solutions was 5134 (A), 2684 (B), 2778 (C), 9189 (D) and 2150 (E). A) Fitting of the IR to the intermediately constricted (65 nm diameter) SpNPC map. All IR homology models were superposed to the integrative model of the single spoke of the IR unit from PDB-DEV_ID 00000051 and the resulting composite model was systematically fitted to the of the individual IR spoke map. B) Fitting of the and NR to the intermediate constricted (65 nm diameter) SpNPC map. The crystal structure with PDB-4GQ2 (covering Nup120-Nup37) and the SpY-complex homology models (covering Nup120-Nup189c-Sec13-Seh1-Nup85-Nup107, Nup107-Nup131/Nup132 and Nup131/Nup132 β-propeller) were superposed with human Y-complexes (PDB-5A9Q) and systematically fitted to the cryo-EM map of an individual NR segment as a single rigid body. C) Fitting of the cytoplasmic side of the intermediately constricted (65 nm diameter) SpNPC map. The inner Y-complex homology model used in (B) was systematically fitted to the cryo-EM map of an individual segment of cytoplasmic side. D-F) same as in (A-C) but fitted to cryo-EM map of the most constricted SpNPC conformation (49 nm diameter).

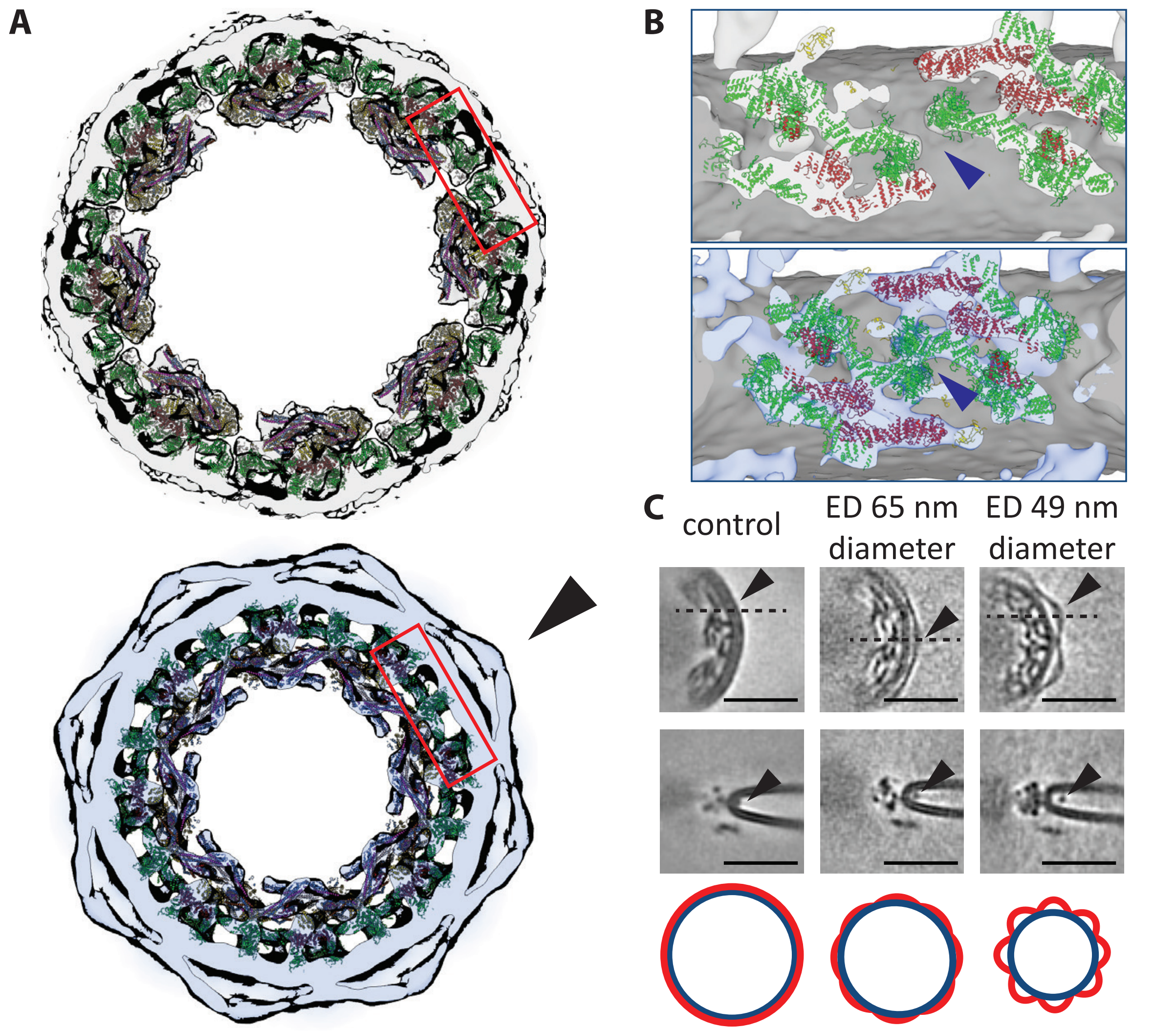


**Supplementary figure 10 Conformational changes of the luminal ring upon NPC constriction.** A) Central slice through actively transporting (top) and the most constricted (bottom) NPC cryo-EM maps superimposed with the respective integrative model (same color code as in **Fig. 1**). Peripheral channels in the area between individual inner ring spokes (indicated in red) are open during active nuclear transport and close during NPC constriction. B) Cutaway side view of the area indicated in (A). Around 3-4 nm wide peripheral gaps are clearly visible between β-propellers of Nup155 (green) from neighbouring IR subcomplexes (blue arrowhead) of actively transporting NPCs (top). The conformational change during NPC constriction leads to closure of the observed gaps in the most constricted NPC state (bottom) through a tight sealing of the now overlapping Nup155 β-propellers potentially affecting transport of inner nuclear membrane proteins (membrane: dark grey; other density: light grey (control) and light blue (ED); Nup models: same color code as **Fig. 1**) C) Luminal densities (black arrowheads, see also A) are arching into the NE lumen during NPC constriction.

Horizontal slices through the inner ring (top) of cryo-EM maps of actively transporting (control; left) intermediate (middle) and most constricted (right) NPCs reveal the appearance of an arch-like density (black arrowheads) during ED, likely corresponding to the C-terminal domains of Pom152, while the N-terminus is attached to the NPC (*49*), which is consistent with previous work on isolated NEs (*34*). Dashed line indicates the position of the orthogonal slices (below) with the luminal density marked by arrowheads. The cartoon (bottom) indicates how a constriction of the nuclear pore membrane (blue) leads to an arching out of the C-terminal luminal domains (red) while the N-terminus remains closely attached to the NE (scale bar: 50nm). In control conditions the luminal densities are directly proximal to the NPC and thus not well discernable form the membrane (scale bars: 50nm).

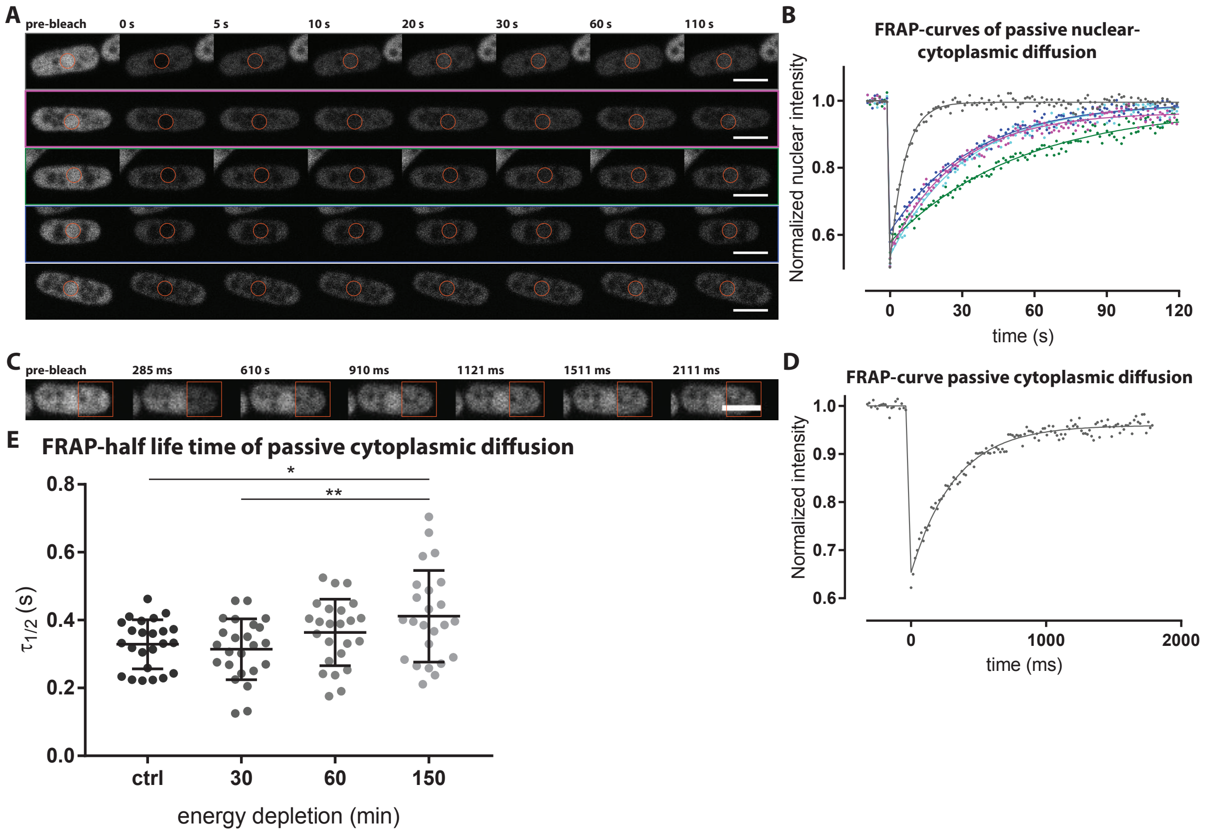


**Supplementary figure 11. FRAP-measurements of passive diffusion across the nuclear envelope and cytoplasmic diffusion.** A) Representative FRAP-images from time series acquired during control conditions (grey) and at 30 min (purple) 60 min (green) 120 min (dark blue) and 210 min (cyan) after ED (from top). The orange circles indicate the bleached area during FRAP-experiments, the time after bleaching is indicated on top (scale bar: 5µm). B) FRAP-curves corresponding to time series in (A). C) Representative snapshots of cytoplasmic diffusion during control conditions. The orange square indicates the area bleached during FRAP-experiment and time stamps indicate time after bleaching (scale bar: 5µm). D) FRAP-curve corresponding to time series in (C). E) Quantification of half-life times extracted from FRAP-curves measuring freely diffusing GFP show only negligible reduction cytoplasmic diffusion during ED with a mean FRAP-recovery half-life time of 0.3288 sec in control conditions and 0.3141 sec 30min, 0.3637 sec 60min, 0.4114 sec 150min after ED. (adjusted p-value: non-significant, <0.05 or <0.01, one-way ordinary ANOVA and Holm-Sidak's multiple comparison test with n=24 for each measurement).

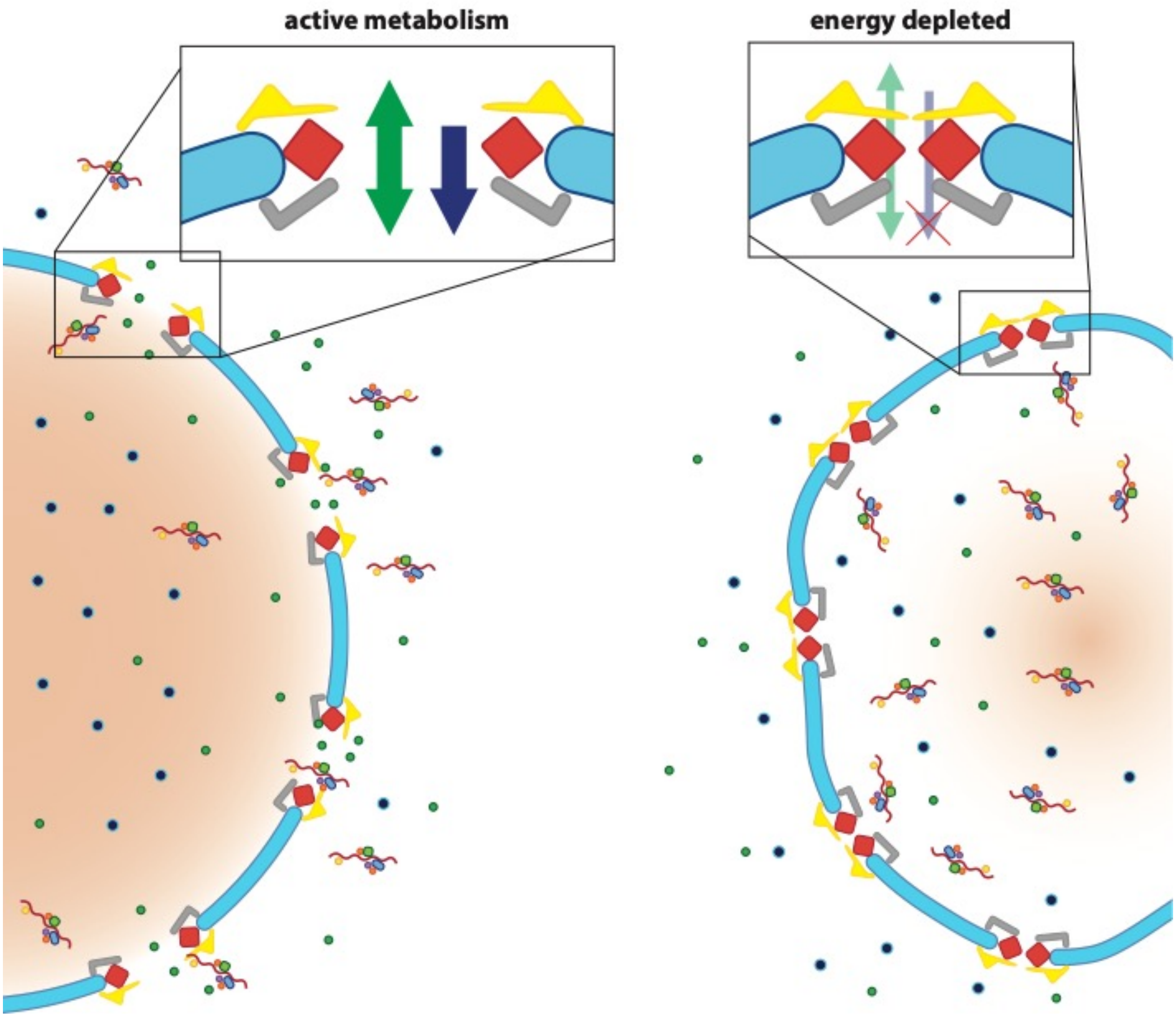


**Supplementary figure 12. A model for NPC constriction during ED.** Concomitant with ED a shrinkage of nuclear volume may lead to a loss of NE tension, thus allowing the NPC to adopt a constricted conformation. During NPC constriction, the central channel volume is reduced to approximately half as compared to metabolically active cells, which likely increases the central channel FG-repeat-domain concentration. A reduced translocation across the NPC of passively diffusing cargoes (green circles; indicated by green arrows in the inserts). According to (*27*), the depletion of GTP results in a decreased RanGTP-RanGDP gradient (orange) and reduced active nuclear import of cargoes containing an NLS (blue circles; indicated by blue arrows in the inserts). According to (*26*), the energy dependent mRNP export is suspended and leads to mRNP accumulation in the nucleus.

| | *S. pombe* | *S. cerevisiae* | *H. sapiens* |
|---|---|---|---|
| **Y-complex** | SpNup85 | ScNup85 | HsNup75/85 |
| | SpNup107 | ScNup84 | HsNup107 |
| | SpEly5 | - | HsELYS |
| | SpNup37 | - | HsNup37 |
| | SpNup120 | ScNup120 | HsNup160 |
| | SpNup131<br>SpNup132 | ScNup133 | HsNup133 |
| | SpNup189C | ScNup145C | HsNup96 |
| | SpSeh1 | ScSeh1 | HsSeh1/Sec13L |
| | - | ScSec13 | HsSec13R |
| | - | - | HsNup43 |
| **Inner ring Nups** | SpNup97/Mug87<br>SpNpp106 | ScNic96 | HsNup93 |
| | SpNup184 | ScNup188 | HsNup188 |
| | SpNup186 | ScNup192 | HsNup205 |
| | SpNup155 | ScNup157<br>ScNup170 | HsNup155 |
| | SpNup40 | ScNup53/59 | HsNup53 |
| | SpNup44 | ScNup57 | HsNup54 |
| | SpNup45 | ScNup49 | HsNup58 |
| | SpNsp1 | ScNsp1 | HsNup62 |
| **Connector Nups** | SpNup189N | ScNup100<br>ScNup116<br>ScNup145N | HsNup98 |
| **Cytoplasmic filament Nups** | - | - | HsNup358 |
| | SpNup82 | ScNup82 | HsNup88 |
| | SpNup146 | ScNup159 | HsNup214 |
| | SpRae1 | ScGle2 | HsRae1 |
| | SpGle1 | ScGle1 | HsGle1 |
| | SpAmo1 | ScNup42/Rip1 | HsNlp1/hCG1/NUPL2 |
| | - | - | HsALADIN |
| | SpDbp5 | ScDbp5 | HsDDX19 |
| **Nuclear basket** | SpNup60 | ScNup60 | - |
| | SpNup61 | ScNup2 | HsNup50 |
| | SpNup124 | ScNup1 | HsNup153 |
| | SpNup211 | ScMlp1<br>ScMlp2 | HsTpr |
| **Trans-membrane Nups** | SpCut11 | ScNdc1 | HsNdc1 |
| | SpPom34/Mug31 | ScPom34 | - |
| | SpPom152 | ScPom152 | HsGP210 |
| | SpTts1 | ScPom33<br>ScPer33 | HsTMEM33 |
| | - | - | HsPom121 |

**Supplementary Table 1. Nucleoporin nomenclature across different species.** Nucleoporin names are indicated for *S. pombe*, *S. cerevisiae* and human, based on (*1*, *93*).

| Strain name | Genotype | Mating type | Source |
|---|---|---|---|
| *wild type* | k972 (*wild type*) | h- | Häring lab |
| *nup37Δ* | nup37::natMX6 | h- | this study |
| ely5Δ | ely5::kanMX leu1-32 ade6-M216 ura4-D18 | h+ | (*43*) |
| nup37Δ-ely5Δ | ely5::kanMX nup37::natMX6 ? ? | h+ | this study |
| GD250 | nup60-mCherry:Kan pBiP1-NLS-GFP-NLS:leu1+_ade-_leu1-32_ura4D-18 | h- | (*66*) |
| AV0890 | ura4+:pact1:sfGFP:terminatortdh1 | h- | (*67*) |
| CZ001 | ura4+:pact1:sfGFP:terminatortdh1 nup60-mCherry:Kann | h- | this study |

**Supplementary Table 2. List of *S. pombe* strains used in this study.**

| average name | resolution | b-factor | No. initial NPCs | No. final particles | No. tomograms | description |
|---|---|---|---|---|---|---|
| *wild type* cytoplasmic side | 22.4 | -2000 | 726 | 2237 | 176 | *wild type* cytoplasmic side |
| *wild type* IR | 23.5 | -2000 | 726 | 1381 | 176 | *wild type* inner ring |
| *wild type* NR | 22.7 | -500 | 726 | 1523 | 176 | *wild type* nuclear ring |
| *nup37Δ* cytoplasmic side | 27.4 | -1000 | 233 | 1064 | 52 | *nup37Δ* knock-out cytoplasmic side |
| *nup37Δ* IR | 26.4 | -1000 | 233 | 1064 | 52 | *nup37Δ* knock-out inner ring |
| *nup37Δ* NR | 30.7 | -1000 | 233 | 1064 | 52 | *Δnup37Δ* knock-out nuclear ring |
| *nup37Δ-ely5Δ* cytoplasmic side | 34.5 | -1000 | 263 | 1217 | 70 | *nup37Δ*-ely5Δ knock-out cytoplasmic side |
| *nup37Δ-ely5Δ* IR | 35 | -2000 | 263 | 1217 | 70 | *nup37Δ*-ely5Δ knock-out inner ring |
| *nup37Δ-ely5Δ* NR | 31.6 | -2000 | 263 | 1217 | 70 | *nup37Δ*-ely5Δ knock-out nuclear ring |
| ED 65 nm diameter cytoplasmic side | 27.4 | Amplitudes matched to defocus data only (see Materials and Methods) | 292 | 1012 | 76 | energy depleted NPCs with initial diameter >50nm cytoplasmic side |
| ED 65 nm diameter IR | 26.9 | Amplitudes matched to defocus data only (see Materials and Methods) | 292 | 1012 | 76 | energy depleted NPCs with initial diameter >50nm inner ring |
| ED 65 nm diameter NR | 28.3 | Amplitudes matched to defocus data only (see Materials and Methods) | 292 | 1012 | 76 | energy depleted NPCs with initial diameter >50nm nuclear ring |
| ED 49 nm diameter cytoplasmic side | 27.8 | Amplitudes matched to defocus data only (see Materials and Methods) | 292 | 533 | 76 | energy depleted NPCs with initial diameter <50nm cytoplasmic side |
| ED 49 nm diameter IR | 26.8 | Amplitudes matched to defocus data only (see Materials and Methods) | 292 | 533 | 76 | energy depleted NPCs with initial diameter <50nm inner ring |
| ED 49 nm diameter NR | 29.7 | Amplitudes matched to defocus data only (see Materials and Methods) | 292 | 533 | 76 | energy depleted NPCs with initial diameter <50nm nuclear ring |

**Supplementary Table 3. List subtomogram averages.** All subtomogram averages are listed with their final resolutions reported according to the 0.143 FSC criterion, the employed b-factor sharpening and the initial and final number of particles and number of used tomograms.

| SpNPC models | SpNPC nup (residue range) | Template structure | Template PDB_ChainName (residue range) | Sequence identity (%) |
|---|---|---|---|---|
| 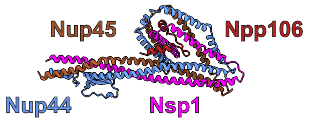 | Nup44 (155-396)<br>Nup45 (239-419)<br>Nsp1 (424-591)<br>npp106 (22-61) | *C. thermophilum* Nup57<br>*C. thermophilum* Nup49<br>*C. thermophilum* Nsp1<br>*C. thermophilum* Nic96 | 5CSW_E (74-314)<br>5CSW_D (244-468)<br>5CSW_C (467-649)<br>5CSW_F (141-180) | 32.0<br>25.4<br>39.5<br>20.5 |
| 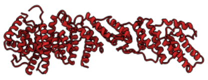 | npp106 (242-929) | *C. thermophilum* Nic96 | 5HB3_C (392-1110) | 26.2 |
| 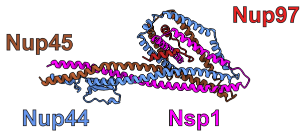 | Nup44 (155-396)<br>Nup45 (239-419)<br>Nsp1 (424-591)<br>Nup97 (12-51) | *C. thermophilum* Nup57<br>*C. thermophilum* Nup49<br>*C. thermophilum* Nsp1<br>*C. thermophilum* Nic96 | 5CSW_E (74-314)<br>5CSW_D (244-468)<br>5CSW_C (467-649)<br>5CSW_F (141-180) | 32.0<br>25.4<br>39.5<br>31.9 |
| 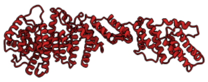 | Nup97 (159-851) | *C. thermophilum* Nic96 | 5HB3_C (392-1110) | 30.1 |
| 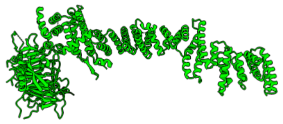 | Nup155 (11-548)<br>Nup155 (549-1315) | *C. thermophilum* Nup170 | 5HAX_A (74-605)<br>5HB1_A (606-1416) | 33.3 |
| 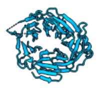 | Nup131 (39-458) | *V. polyspora* Nup133 | 4Q9T_A (63-497) | 20.3 |
| 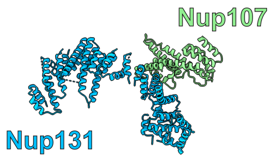 | Nup131 (473-927)<br>Nup131 (928-1142)<br>Nup107 (580-813) | *H. sapiens* Nup133<br>*S. cerevisiae* Nup133<br>*H. sapiens* Nup107 | 3I4R_B (518-1003)<br>3KFO_A (956-1142)<br>3I4R_A (667-924) | 20.1<br>17.9<br>20.9 |
| 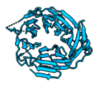 | Nup132 (37-446) | *V. polyspora* Nup133 | 4Q9T_A (63-497) | 19.9 |
| 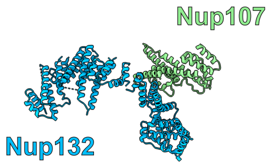 | Nup132 (459-933)<br>Nup132 (934-1162)<br>Nup107 (580-813) | *H. sapiens* Nup133<br>*S. cerevisiae* Nup133<br>*H. sapiens* Nup107 | 3I4R_B (518-1003)<br>3KFO_A (956-1142)<br>3I4R_A (667-924) | 20.1<br>17.9<br>20.9 |
| 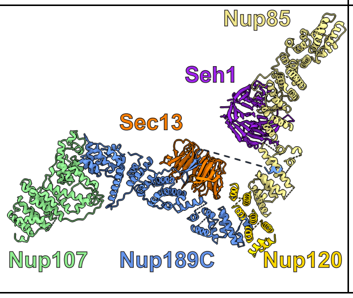 | Nup107 (73-512)<br>Nup189C (189-841)<br>Sec13 (8-293)<br>Seh1 (6-331)<br>Nup85 (8-671)<br>Nup120 (954-1115) | *S. cerevisiae* Nup84<br>*S. cerevisiae* Nup145<br>*S. cerevisiae* Sec13<br>*S. cerevisiae* Seh1<br>*S. cerevisiae* Nup85<br>*S. cerevisiae* Nup120 | 4XMM_F (7-442)<br>4XMM_B (92-712)<br>4XMM_A (8-293)<br>4XMM_C (1-346)<br>4XMM_D (47-744)<br>4XMM_E (884-1036) | 22.8<br>23.9<br>59.2<br>44.3<br>19.9<br>20.4 |

| | | | | |
|---|---|---|---|---|
| 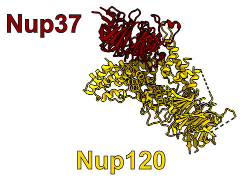 | Nup120 (1-949) | *S. pombe* Nup120 | 4GQ2_M (1-949) | not applicable (crystal structure) |
| | Nup37 (1-387) | *S. pombe* Nup37 | 4GQ2_P (1-387) | not applicable (crystal structure) |
| 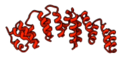 | Ely5 (88-298) | *S. pombe* Nup120 | 4FHN_B (610-985) | 10.1 |
| 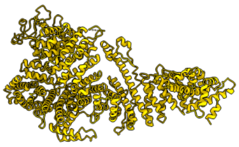 | Nup184 (1-970) | *M. thermophila* Nup188 | 4KF7_A (19-1149) | 17.2 |
| | Nup184 (971-1439) | *C. thermophilum* Nup192 | 5HB4_B (1045-1598) | 15.6 |
| | Nup184 (1440-1564) | *M. thermophila* Nup188 | 4KF8_A (1623-1769) | 18.4 |
| 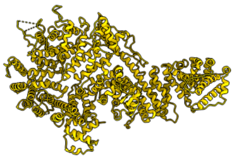 | Nup186 (1-336) | *C. thermophilum* Nup192 | 4KNH_A (1-352) | 18.4 |
| | Nup186 (337-1647) | | 5HB4_A (353-1728) | |

**Supplementary Table 4. List of *S. pombe* homology model.** The homology modeled Nucleoporins and the corresponding templates are listed.

**Supplementary video 1. Overview of the integrative model of the actively transporting SpNPC**. The structural model is shown as cartoon representation, colored by Nup identity as in **Fig. 3**, and shown fitted to the cryo-EM map (grey).

**Supplementary video 2. Overview of the integrative model of SpNPC upon ED with diameter of 65 nm**. The structural model is shown in cartoon representation, colored by Nup identity as in **Fig. 3**, and shown fitted to the cryo-EM map (grey).

**Supplementary video 3. Overview of the integrative model of SpNPC upon ED with diameter of 49 nm**. The structural model is shown in cartoon representation, colored by Nup identity as in **Fig. 3**, and shown fitted to the cryo-EM map (grey).

**Supplementary video 4. Morph between the SpNPC conformations at three different diameters as seen from the cytoplasmic side.** The structural models are shown in cartoon representation, colored by Nup identity as in **Fig. 3**. The morph was calculated using UCSF ChimeraX. Please note that the morph helps visualizing the conformational differences but does not necessarily depicts the realistic path of the conformational changes during dilation and constriction.

**Supplementary video 5. Morph between the SpNPC conformations at three different diameters as seen from the nuclear side.** The structural models are shown in cartoon representation, colored by Nup identity as in **Fig. 3**. The morph was calculated using UCSF ChimeraX. Please note that the morph helps visualizing the conformational differences but does not necessarily depicts the realistic path of the conformational changes during dilation and constriction.

**Supplementary video 6. Morph between the SpNPC conformations at three different diameters in side view.** The structural models are shown in cartoon representation, colored by Nup identity as in **Fig. 3**. The morph was calculated using UCSF ChimeraX. Please note that the morph helps visualizing the conformational differences but does not necessarily depicts the realistic path of the conformational changes during dilation and constriction.

**Supplementary video 7. Morph between the SpNPC conformations at three different diameters as cutaway view.** The structural models are shown in cartoon representation, colored by Nup identity as in **Fig. 3**. The morph was calculated using UCSF ChimeraX. Please note that the morph helps visualizing the conformational differences but does not necessarily depicts the realistic path of the conformational changes during dilation and constriction.
